# Swinging lever mechanism of myosin directly demonstrated by time-resolved cryoEM

**DOI:** 10.1101/2024.01.05.574365

**Authors:** David P. Klebl, Sean N. McMillan, Cristina Risi, Eva Forgacs, Betty Virok, Jennifer L. Atherton, Michele Stofella, Donald A. Winkelmann, Frank Sobott, Vitold E. Galkin, Peter J. Knight, Stephen P. Muench, Charlotte A. Scarff, Howard D. White

## Abstract

Myosins are essential for producing force and movement in cells through their interactions with F-actin. Generation of movement is proposed to occur through structural changes within the myosin motor domain, fuelled by ATP hydrolysis, that are amplified by a lever swing^1^, transitioning myosin from a primed (pre-powerstroke) state to a post-powerstroke state. However, the initial, primed actomyosin state, proposed to form prior to lever swing, has never been observed. Nor has the mechanism by which actin catalyses myosin ATPase activity been resolved. To address this, we performed time-resolved cryoEM of a myosin-5 mutant having slow hydrolysis product release. Primed actomyosin was captured 10 ms after mixing primed myosin with F-actin, whereas post-powerstroke actomyosin predominated at 120 ms, with no abundant intermediate structures. The structures were solved to 4.4Å and 4.2Å global resolution respectively. The primed motor binds to actin through its lower 50 kDa subdomain, with the actin-binding cleft open and Pi release prohibited. N-terminal actin interactions with myosin promote rotation of the upper 50 kDa subdomain, which closes the actin-binding cleft, and enables Pi release. Formation of upper 50 kDa subdomain interactions with actin creates the strong-binding interface required for effective force production. The myosin-5 lever swings through an angle of 93°, predominantly along the actin axis, with little twisting, to produce the post-powerstroke state. The magnitude of the lever swing matches the typical step length of myosin-5 walking along actin. These time-resolved structures directly demonstrate the swinging lever mechanism, ending decades of conjecture on how myosin produces force and movement.

## Main text

Myosins are molecular motors that move or move along filamentous actin (F-actin). They perform many functions in eukaryotes, ranging from muscle contraction to organelle transport, with mutations linked to a range of diseases including heart disease, deafness and cancer^2^. Myosins comprise a motor domain, which can be divided into 4 subdomains (N-terminal, upper 50 kDa (U50), lower 50 kDa (L50) and converter), a light-chain binding domain and a tail region. The converter and light-chain binding domain form the lever, that rectifies and amplifies changes within the motor domain^3^.

ATP hydrolysis by myosin provides the energy for doing work. In the nucleotide-free state, the myosin motor is strongly bound to F-actin^4,5^. ATP-binding opens a cleft between the U50 and L50 domains, reducing the affinity of myosin for F-actin, which dissociates the complex^6^. Once detached, myosin undergoes the recovery stroke, in which the myosin lever becomes primed to generate force, followed by ATP hydrolysis to ADP and phosphate (Pi)^7^. Release of the Pi from myosin is slow, precedes release of ADP, and thus limits the rate of energy release in the absence of interactions with actin. Primed myosin, with ADP and P_i_ bound, rebinds F-actin leading to P_i_-release, cleft closure and generation of movement, proposedly through swinging of the lever (powerstroke)^8,9^, towards the barbed-end of F-actin (+actin)^9^ for the majority of myosins. Actin accelerates Pi release ∼1000-fold. The order in which Pi-release, cleft closure and powerstroke occur is debated ^10,11^. Release of ADP from the complex is, in some myosins, coupled to a second, smaller swing of the lever that completes the structural cycle^12,13^.

The mechanisms of force generation and actin activation of ATPase activity remain controversial, in part due to a lack of structural information on how myosin initially interacts with actin in its primed state^3,14^. Actomyosin structures in the ADP and nucleotide-free states, obtained by cryo electron microscopy (cryoEM), reveal the architecture of strongly-bound actomyosin complexes in which both the U50 and L50 subdomains interact with actin, the cleft is closed, and the lever adopts a post-powerstroke (postPS) position^15^. The structure of the myosin motor in the primed state in the absence of actin, with ADP-P_i_ or analogues in the nucleotide binding site, has been solved by X-ray crystallography for multiple myosin classes including myosin-2^16^,-5^12^ and -6^17^. The myosin primed state structures show an open cleft between the U50 and L50 subdomains, and a primed lever^12^. However, previous attempts to image its attachment to actin have failed.

At steady state, attached primed myosin is rare because it is a weakly-bound state that rapidly transitions to a postPS strongly-bound state. Thus, the traditional high-resolution structural methods, X-ray crystallography and cryoEM plunge-freezing approaches, are unable to capture a primed actomyosin structure. Here, we have overcome these difficulties by using a myosin-5 mutant construct with higher affinity for actin^18^ and an increased lifetime of the attached primed state^19^, and by using a microspray method for cryoEM specimen preparation^20^ that permits millisecond time resolution. We present a structure of primed actomyosin at 4.4 Å global resolution, which shows how the myosin motor interacts with F-actin in its primed state to initiate force generation and directly demonstrate the swinging lever mechanism.

### Trapping primed actomyosin by time-resolved cryoEM

To trap the actomyosin primed complex, we pre-incubated a myosin-5 construct (motor domain plus 1 IQ light-chain binding domain) with ATP for ∼2s, allowing the myosin to bind and hydrolyse ATP, so it was primed for actin binding^19^. This was then mixed rapidly with F-actin, sprayed onto an EM grid, and plunge-frozen to trap the reaction after 10 or 120 ms using our custom-built device (see Methods and Extended Data Fig. 1)^20,21^. We used a myosin-5 mutant with an S^217^A mutation in switch 1 in the nucleotide binding pocket and DDEK^594-597^ deletion in loop 2 (Extended Data Fig. 2). S^217^A slows actin-activated Pi release (198 s^-1^ to 16 s^-1^)^19^ and the deletion increases the affinity of the myosin-5-ADP-Pi primed state for F-actin ∼10 fold^18^. This double mutant motor is fully functional in actin-motility assays and has a maximum actin-activated Pi-release rate of 13 s^-1^ (Extended Data Fig. 3).

We chose two timepoints at which to vitrify the myosin-actin mixture, 10 and 120 ms. At 10 ms, the maximum speed of the setup, based on the kinetic data, we expected the majority of actomyosin complexes to be in the primed state, whereas at 120 ms, a higher proportion of these would have transitioned to a postPS state, ensuring that any intermediate states between the primed and postPS could be captured (see Extended Data Fig. 3).

The time-resolved cryoEM data yielded two distinct classes of actomyosin-5 structures, which we identified as the primed and postPS states, and solved to global resolutions of 4.4 and 4.2 Å, respectively (Extended Data Fig. 4). CryoEM density maps were fitted with atomic models to enable detailed interpretation, complemented by molecular dynamics simulations (see Methods, and Extended Data Table 1 & 2). Calmodulin density in all the EM maps is weak, indicating low occupancy of the heavy chain by calmodulin. The postPS actomyosin structure was similar to previous structures of strongly-bound states^15^.

The lever swing mechanism predicts that upon mixing of primed myosin with F-actin, primed actomyosin will initially predominate with postPS actomyosin accumulating over time. We found that 62 % of actomyosin complexes were in the primed state at 10 ms (Extended Data Fig. 5). At 120 ms, the proportion of primed actomyosin complexes was reduced to 36 % concomitant with an increase in postPS complexes, in reasonable agreement with a Pi-release rate of 13 s^-1^ (Extended Data Fig. 3d). Intermediate states were not detected despite extensive 3D classification and masking (Extended Data Fig. 4). This time-dependence of conformation directly demonstrates the swinging lever mechanism.

### Structure of actomyosin in the primed state

In the primed state, myosin interacts with actin through its L50 domain (Fig. 1, Supplementary Video 1). The central actomyosin interface is formed between two neighbouring actin subunits and the myosin helix-loop-helix (HLH) motif (Fig. 1a-e), with additional interactions between actin and myosin loop2 and myosin loop3 completing the interface (Fig. 1f, g). The main contacts are primarily hydrophobic in nature, supplemented by electrostatic interactions. The HLH-actin interactions are the same as observed for the strong-binding states^22^, largely conserved across myosin classes in higher eukaryotes^23^. Thus, the orientation of the primed motor domain when docked onto actin resembles that of strongly-bound states except that the U50 does not interact with actin in primed actomyosin (Fig. 1b). Residues at the tip of the HLH loop (M515, P516) fit into a hydrophobic pocket on actin created by conserved residues in the pointed-end F-actin (-actin) subdomain-1 (I345-L349) and subdomain-3 (A144), and the +actin subdomain-2 D-loop (M44-M47). Residues E511, K514 and K517 in the HLH motif can form hydrogen bonds with S350/T351, S348, and G146 backbone respectively (Fig. 1e).

**Figure 1.**
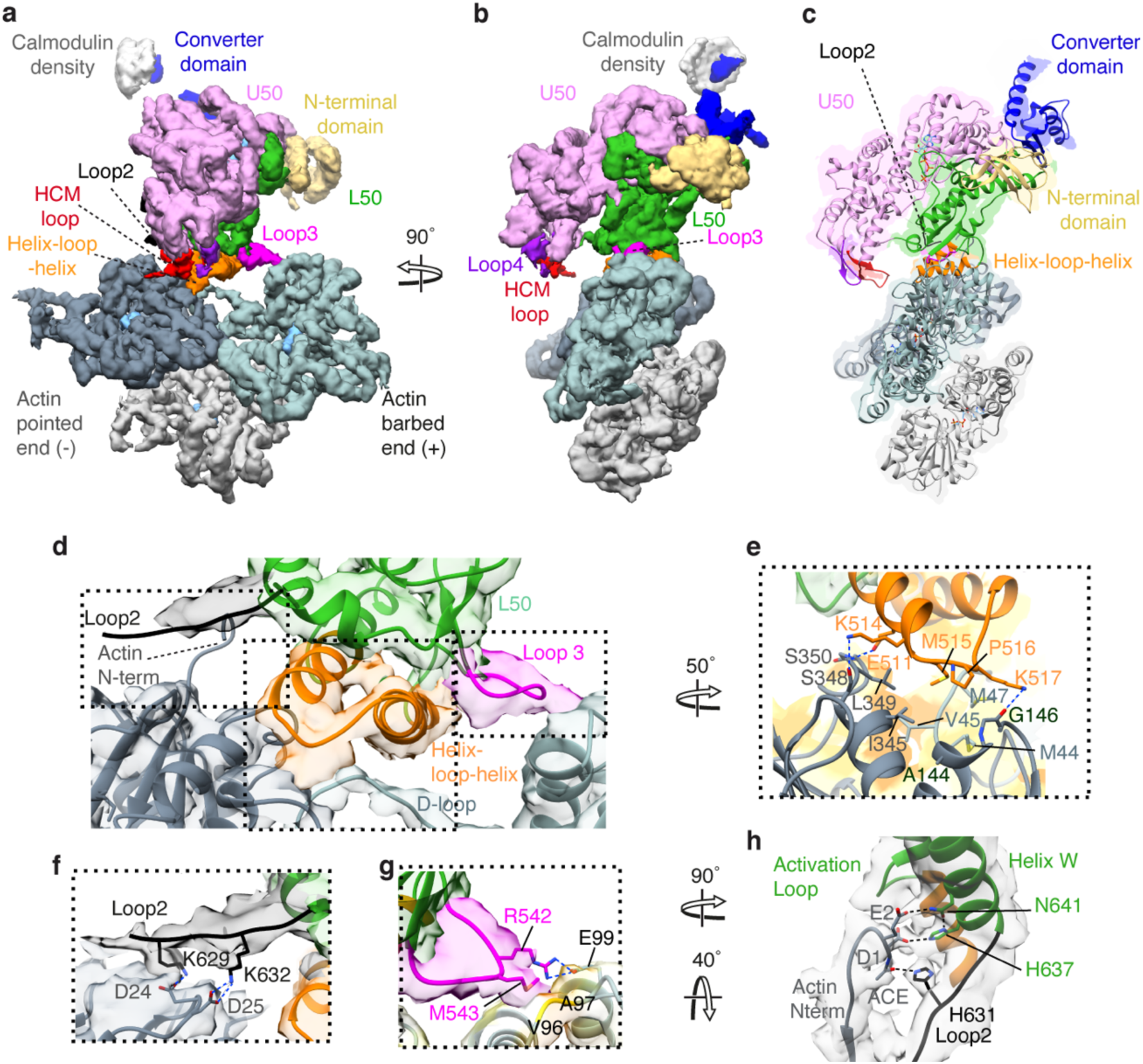
Structure of the primed actomyosin-5 complex. (a, b) CryoEM density map of the primed actomyosin-5 complex, segmented and coloured by myosin subdomains and actin chains as indicated (with central three actin subunits displayed). Actin subunits are shown in slate grey(-end), blue-grey (+end), and light grey. Map thresholded to show secondary structure (myosin 0.085, actin 0.2) and shown (a) in side view of F-actin and (b) in end-on view of F-actin, looking towards the pointed end. (c) Backbone depiction of atomic model of primed actomyosin-5, fitted into the EM density map, viewed as in (b). (d) Magnified side view of the actomyosin interface, contacts are made by (e) the myosin HLH motif, (f) loop2 (threshold 0.007*) and (g) loop3. Relevant interacting residues are labelled and shown. In (e) and (g), HLH and actin segmented maps are coloured by hydrophobicity (orange most hydrophobic to white hydrophilic) to highlight hydrophobic interactions, especially the hydrophobic pocket formed by the two neighbouring actin subunits into which the tip of the HLH motif fits. (h) Magnified view showing the N-terminal residues of the -actin subunit (slate grey), side chains of D1, E2 and acetyl group (ACE) of D1, reaching out to interact with Helix W of the L50 domain, and loop2, at H637 and N641, and H631 respectively (EM density threshold 0.007*). DeepEMhancer post-processed map depicted in (a-e, g), and *RELION post-processed map in (f,h).

Myosin loop2 is flexible and poorly resolved in the primed state, as in most other actomyosin structures^15,22^. Yet, the C-terminal portion of loop2 (residues 628-632) has appreciable density that adopts an elongated conformation, reaching out parallel to the actin surface, allowing positively charged residues K629 and K632 to interact with the negatively charged D24 and D25 in -actin subdomain-1, respectively (Fig. 1f). A ridge of weaker density extends further along the surface of the actin suggesting that more of loop2 may be associated with the actin surface. In myosin loop3, M543 can interact hydrophobically with residues V96 and A97 of +actin subdomain-1, enabling R542 to form an ionic interaction with E99 of +actin subdomain-1 (Fig. 1g).

The converter is in a primed position within the motor domain, and the orientation of the motor domain on actin results in the emerging lever helix pointing along the actin axis towards the pointed end, at an angle of ∼52° to the actin axis.

The N-terminal residues of actin (residues 1-4, DEDE), which are unresolved in most actin structures, reach out to interact with Helix W of the myosin L50 subdomain and loop2. Actin residues D1 and E2 interact with H637 and N641 in helix W respectively, and the acetyl group of the acetylated N-terminal residue D1 interacts with H631 in loop2 (Fig. 1h). These N-terminal actin interactions with myosin lead to subtle changes in primed myosin structure, described below, that may suggest how actin activates myosin ATPase activity.

### Structural changes in primed myosin upon actin binding

The time-resolved cryoEM data contained unbound myosin-5 molecules providing us with the opportunity to directly compare myosin structure in the unbound and actomyosin states (Fig. 2 and Supplementary Video 2). Unbound myosin motors from the 120 ms data were analysed to produce an EM map with global resolution of 4.9 Å (Extended Data Fig. 6). This revealed that unbound myosin motors were in a primed state, vitrified prior to productive actin binding. The crystal structure of the myosin-5c motor domain trapped in the primed state by use of ADP-vanadate (PDB ID: 4ZG4) was well accommodated within the cryoEM density^24^, except in the position of the converter domain and relay helix (Extended Data Fig. 7a-e). Thus, flexible fitting of the crystal structure in the map was used to produce a model of our myosin-5a construct in the unbound primed state (Extended Data Fig. 7a,c,e).

**Figure 2:**
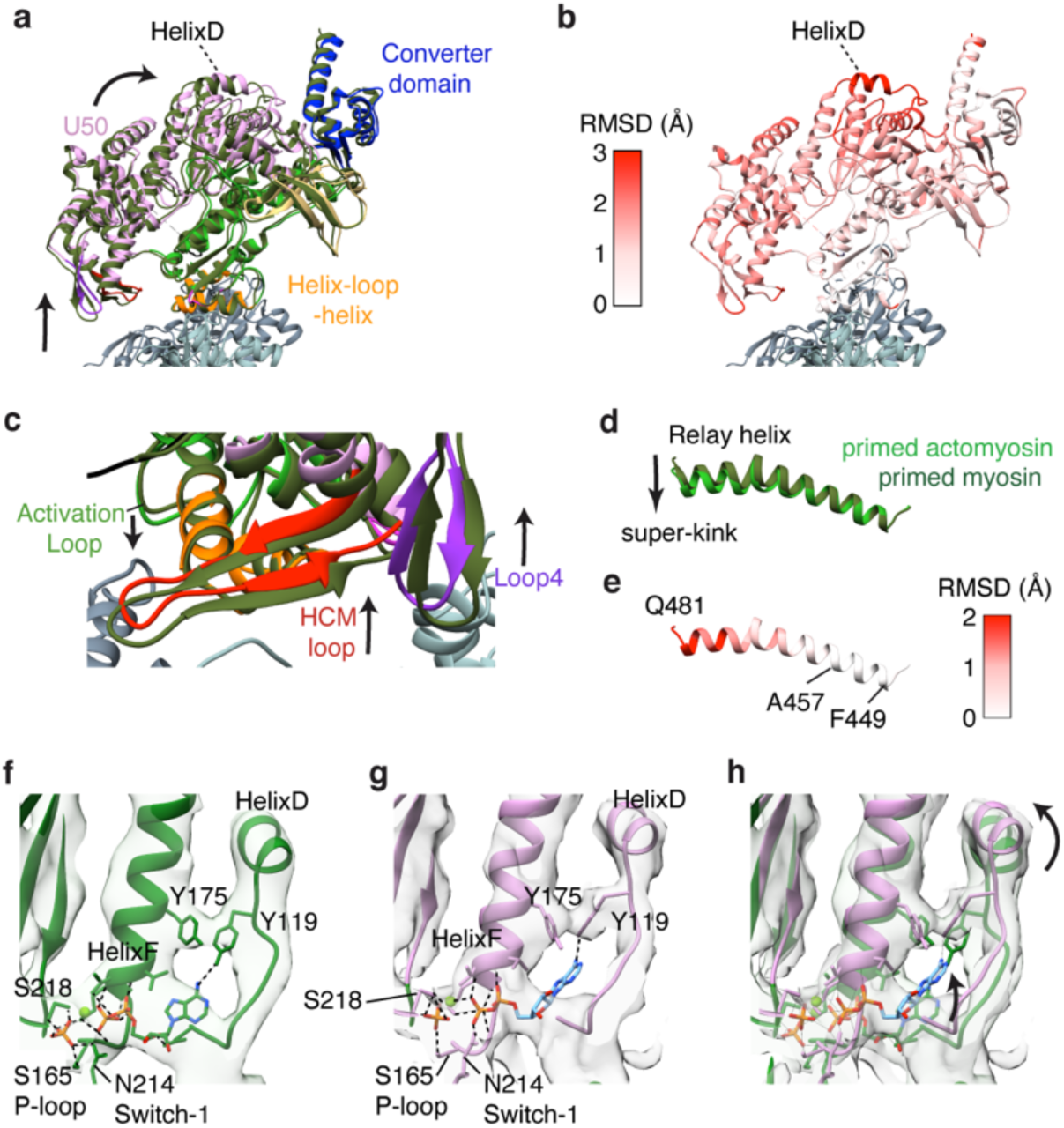
Comparison of myosin structure in the primed actomyosin complex with unbound primed myosin. (a) Superposition of the primed actomyosin (coloured as in Fig. 1) and unbound primed myosin (forest green) aligned on the core primed actomyosin interface (HLH motif, residues 505-530). View towards actin pointed end. (b) Corresponding RMSD of myosin residues between primed actomyosin and primed myosin, showing greatest movement occurs in HelixD. The whole U50 is rocked back, around the actin axis, towards the converter domain, resulting in (c) the HCM loop and loop4 moving away from the actin surface. The activation loop also extends down, reaching out to the actin surface. (d) The myosin relay helix, when aligned on residues 449-457 at the start of the relay helix, is found to have additional kinking upon actin binding. (e) RMSD of primed actomyosin relay helix relative to primed myosin. (f) Unbound primed myosin and (g) primed actomyosin models focussed on HelixD, Y119, Y175 and nucleotide, overlaid with their respective cryoEM maps, thresholded equivalently. (h) Overlay of (f) and (g) showing movement of HelixD upon binding of myosin to actin causes rearrangement of tyrosine residues Y119 and Y175, resulting in larger freedom of placement of ADP in the nucleotide binding pocket. The Pi is anchored by interactions with P-loop/HelixF (S165/K169) and Switch-1 (N214/S218). Primed actomyosin RELION post-processed map depicted throughout.

The myosin models for unbound primed myosin-5 and primed actomyosin-5 are very similar (0.80 Å RMSD from global alignment of the motor domains across 708 Cα atom pairs) yet subtle changes are seen in the flexible regions, especially in the position of the converter domain and HelixD (Extended Data Fig. 7f,g). When the two structures are aligned on the main actomyosin binding interface, the HLH (residues 505-530) alone (Fig. 2a), the entire U50 is observed to be displaced with the largest shift in the position of HelixD (Fig. 2b). This suggests that subtle structural changes in the myosin motor are induced by actin binding and propagated through the molecule.

In the bound state, with myosin anchored to actin through the HLH, the rest of the L50 moves downwards, relative to the actin axis, so that the U50 domain is rotated circumferentially around F-actin towards the converter, which lifts the HCM loop and loop4 further away from the actin surface (Fig. 2a,c, Supplementary Video 2). The interaction of the N-terminal residues (1-2) of actin with the neighbouring HelixW and loop2 (Fig. 1h) could drive this motion. The C-terminal end of loop2 extends to contact the actin surface (Fig. 1e), and the ‘activation loop’ (residues 501-504, between HelixQ and HelixR) protrudes further out from the axis of its neighbouring helices, reaching out for the actin surface (Fig. 2c). There is an increase in the bend of the relay helix when primed myosin binds to actin suggesting it is under additional strain (Fig. 2d, e). Yet interestingly, the converter hardly moves relative to the HLH motif (Fig. 2b), such that there is little movement of the lever when primed myosin binds to actin.

The movement of HelixD results in the rearrangement of the position of Y175 and Y119 (Fig 2. f,g,h and see Supplemental Video 2). These residues interact with the adenosine ring of the nucleotide and the rearrangement likely results in less restraint on ADP in the nucleotide pocket. This is supported by the observation of weaker density for the adenosine ring in the primed actomyosin EM density in comparison to that in the unbound primed myosin EM density (Fig 2. f,g). Consequently, in the primed actomyosin model, refined with molecular dynamics in ISOLDE^25^, the ADP is placed further back into the pocket, towards HelixD, creating strain that may promote Pi dissociation from the ADP moiety since the Pi is anchored by interactions with the P-loop/HelixF (S165 and K169 respectively) and Switch-1 (N214 and S218) (Fig 2. f,g,h). However, Pi cannot dissociate because its exit route is blocked as the salt bridge between R219 in switch-1 and E442 in switch-2, termed the backdoor, is still intact.

### Structural changes during the power stroke

Comparison of the actomyosin primed and postPS states allows us to describe the structural changes that occur during the power stroke. The biggest change is the large-scale movement of the converter and light-chain binding domain (Fig. 3a-e, Extended Data Fig. 8 and Supplementary Video 3), responsible for generation of external mechanical force. The lever swings through ∼93°, predominantly along the actin axis, and is displaced azimuthally by only 4° right-handed (Fig. 3c-e), with a small (2.5°) right-handed torsion of the lever around its own axis. The N-terminal domain is displaced by ∼10 Å orthogonally to the actin axis (Fig. 3e). Thus, the myosin-5 motor successfully converts complex internal movements into a simple swinging motion along actin and these data directly demonstrates the swinging lever mechanism.

**Figure 3:**
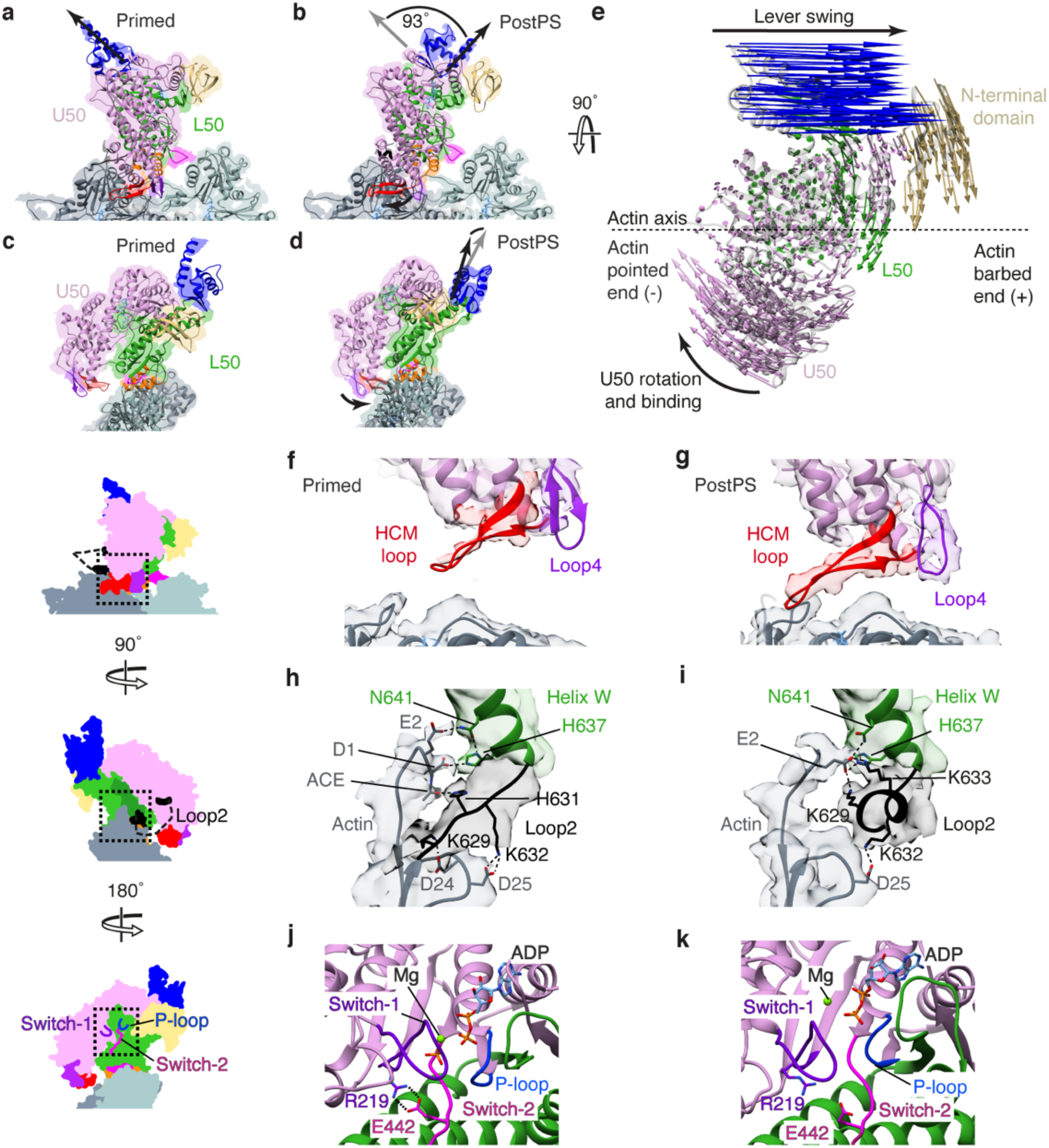
Structural changes during the power stroke. (a) Primed actomyosin structure (as shown in Fig. 1a) and (b) corresponding view of the postPS actomyosin structure with lever positions indicated by a black arrow. The lever swings ∼93° between structures, in a slight right-handed arch (4°). (c) In end-on view, we observe that primed actomyosin has an open actin-binding cleft, while (d) postPS actomyosin has a closed cleft. (e) In aerial view, vectors depict the movement of myosin residue Cα atoms between primed and postPS actomyosin states. The biggest motions are attributable to lever swing, U50 rotation and binding to actin, and displacement of the N-terminal domain. The HCM-loop and loop4 are distant from the actin surface in the (f) primed state but interact with actin in the (g) postPS state, EM density segmented and coloured by myosin subdomains (contour level 0.008). N-terminal actin interactions with loop2 and helixW are changed between (h) primed and (i) postPS states. Nucleotide binding site in (j) primed and (k) postPS structures. The ‘backdoor’ (salt bridge between R219 and E442) is opened through rotation of the U50 and switch-1 and P-loop moving away from switch-2. DeepEMhancer post-processed map depicted in (a-d, f, g), and *RELION post-processed map in (h, i).

Whilst the interactions between actin and the myosin L50 domain (HLH motif and loop3) remain largely unchanged between the primed and postPS state, the U50 interactions are distinctly different (Fig. 3f, g). In the primed state, the HCM loop and loop4 are poorly resolved, indicating flexibility in this region, and both loops are too distant from the actin surface to form stable contacts with it (Fig. 3f). In the postPS state the U50 domain is rotated such that the actin-binding cleft is closed and the HCM loop and loop4 can interact with the actin surface, forming both hydrophobic and charged interactions (Fig. 3g & Extended Data Fig. 9a-d), as seen in previous strongly-bound actomyosin structures^15,22,23^. These additional interactions increase the surface area of the binding interface from 375 Å^2^ in the primed state to 729 Å^2^ in the postPS state, creating a much stronger binding interface and providing the structural basis for the weak to strong binding transition.

In the postPS actomyosin structure, the interactions of loop2 with -actin subdomain 1 are different to those seen in the primed structure (Fig 3h, i). The interactions of H631 with acetyl-D1 and K629 with D24 (Fig. 3h) are broken and the C-terminal portion of loop2 adopts a helical conformation with K629 and K633 forming stronger ionic interactions with actin E2 (Fig. 3i). The preserved interaction of K632 in loop2 with D25 means that the change in loop2 conformation, which shortens loop2, would rotate the U50 around towards the actin surface, resulting in formation of the second binding interface and cleft closure (see Supplementary Video 3). An interaction between residue K502 in the activation loop and E4 of actin is also formed in the postPS state (Extended Data Fig. 9e).

Within the nucleotide binding pocket, there are relative movements between switch-2, switch-1 and the P-loop that indicate that Pi has been released in the postPS structure (Fig. 3j, k). The salt bridge between R219 in switch-1 and E442 in switch-2 (termed the backdoor) is intact in the primed actomyosin structure and broken in the postPS structure. Rotation of the U50 across the L50, resulting in cleft closure, displaces switch-1 and the P-loop away from switch-2 to open the backdoor and enable Pi release (see morph between primed and postPS actomyosin in Supplemental Video 3).

Our postPS actomyosin-5 structure shows a closed actin-binding cleft as well as a postPS lever position (Fig. 3d) and has high similarity to previous structures of strongly-bound actomyosin complexes (ADP-bound or rigor states)^15^. The cryoEM density shows clear evidence for the presence of MgADP (Extended Data Fig. 9f) and we, therefore, identify the postPS state as ADP-bound actomyosin-5. We observe that the position of the lever is more similar to that observed in previous rigor structures (Extended Data Fig. 9g), rather than ADP-bound structures^12,15^. We also observe that the density for the magnesium ion in the nucleotide binding pocket (Extended Data Fig. 9h, i) is in a different position to that seen in other ADP-bound structures^15^. This could be due to the S217A mutation changing the Mg coordination, and may explain the 2-fold increase in ADP release rate for the S217A mutant compared to WT^19^, along with the change in lever position.

### Structural mechanism of myosin force generation and ATPase activation on F-actin

The changes we observe between unbound primed myosin, primed actomyosin and postPS actomyosin allow us to propose the mechanism by which myosin generates movement and actin catalyses it (Fig. 4, Supplementary Video 4).

**Figure 4:**
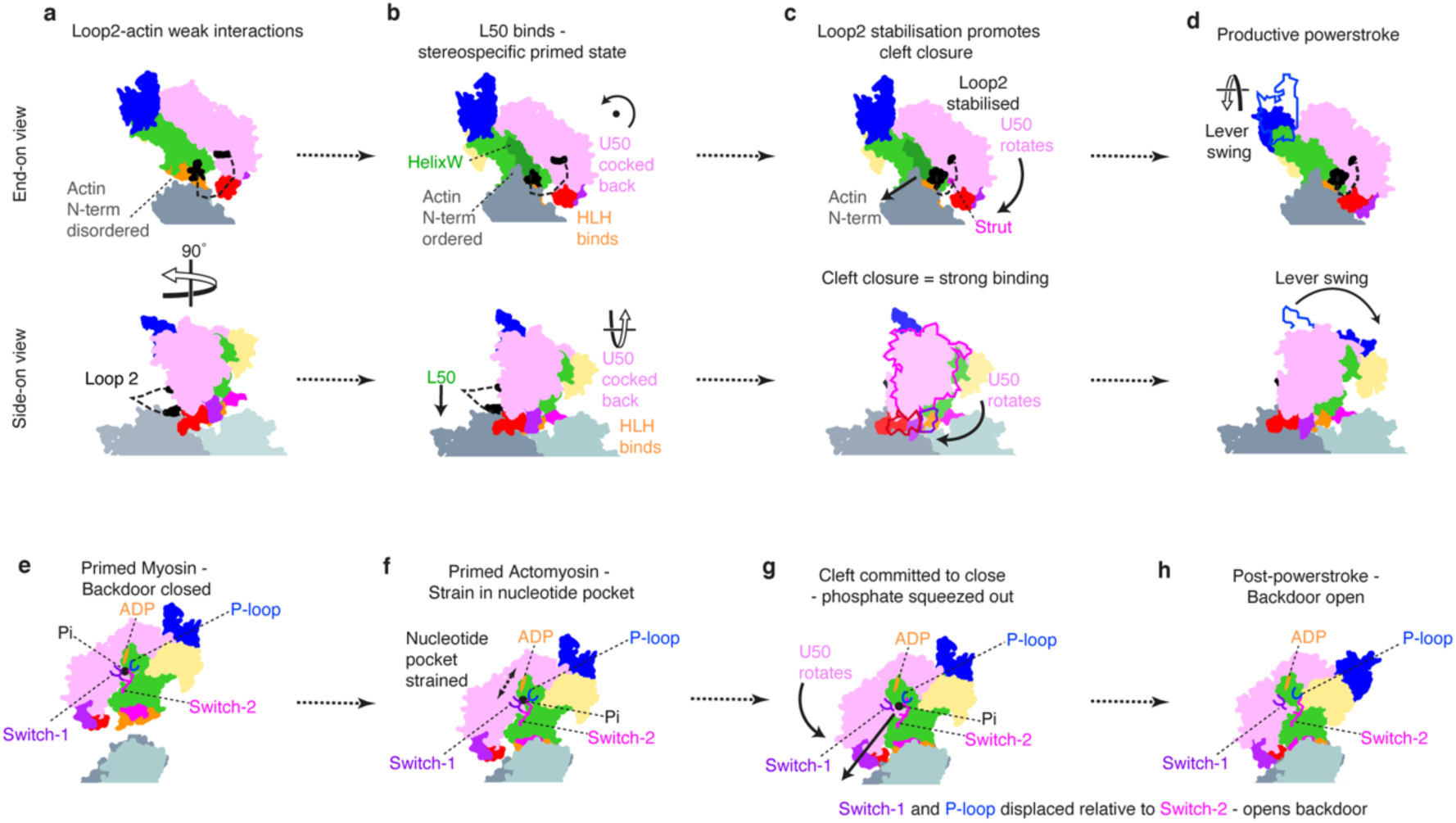
Models of myosin force generation and ATPase activation on F-actin. (a-d) force generation, upper row: end-on view; lower row: side view. (e-h) ATPase activation (a) Primed myosin initially binds weakly to actin through electrostatic interactions of loop2 with actin subdomain-1. This brings the L50 of myosin in close proximity to the actin surface, enabling formation of the stereospecific primed actomyosin state. (b) HLH binding enables the actin N-terminal residues 1-2 to interact with HelixW and loop2, resulting in the U50 being cocked back towards the converter domain, rotated around the F-actin axis. (c) Rearrangement of N-terminal actin interactions with HelixW and loop2 result in loop2 stabilisation at its C-terminal end. This shortens loop2, rotating the U50 and attracting the negatively-charged strut to positively-charged loop2, promoting cleft closure. (d) Cleft closure results in the strong binding interface needed to sustain force and concomitantly results in twisting of the transducer, straightening of the relay helix and lever swing. (e) In the unbound primed state, the backdoor is closed, prohibiting Pi release. (f) Upon binding of primed myosin to actin, cocking back of the U50 towards the converter creates strain in the nucleotide pocket, with the ADP drawn away from the well-coordinated Pi, prohibiting reversal of hydrolysis and promoting Pi release. (g) As the U50 rotates, and the initial interactions between the U50 and the actin surface are formed, switch-1 and the P-loop are displaced relative to switch-2, the backdoor is opened, and Pi is squeezed out into the Pi release tunnel. (h) In the PostPS state, Pi has been released and the lever has swung and the backdoor is open. Pi re-entry into the nucleotide pocket is highly unfavourable.

It is generally accepted that myosin initially binds weakly to actin through interactions between positively-charged residues of loop2 and negatively-charged residues in actin subdomain-1 (Fig. 4a)^22^, which are indeed seen in our primed structure (Fig1f). This brings the L50 in close proximity to the actin surface, enabling the stereospecific interaction between the L50 (HLH and loop3) and F-actin to form quickly after this initial interaction. This interaction triggers a significant rearrangement within the primed myosin (cocking back of the U50) to produce the primed actomyosin we observe (Fig. 4b). In the transition between primed and postPS states, we show that the U50 must rotate resulting in cleft closure and producing the strong binding interface required to sustain the force generated by lever swing (Fig. 4c-d). Yet, the question of how actin activates myosin ATPase activity still remains.

Actin N-terminal residues 1-4 are implicated in myosin ATPase activation, as deletion or mutation of these residues diminishes actin-activated ATPase activity^26,27^. We find that actin structure is almost unchanged between free actin, primed actomyosin and postPS actomyosin, except in the N-terminal residues, which are disordered in free actin, become ordered in primed actomyosin and adopt a different conformation in postPS actomyosin (Extended Data Fig. 10). Density for the D-loop in subdomain-2 is also stronger in the actomyosin structures in comparison to free actin due to stabilisation upon myosin binding. When primed myosin binds to F-actin, actin residues 1-2 interact with both HelixW and loop2. The interactions with HelixW provide stabilisation of the L50 and cause a slight rotation of the U50 back towards the converter domain (Fig. 4b), which results in HelixD movement creating strain in the nucleotide binding pocket that would enable Pi dissociation, yet Pi cannot dissociate because the back door is still closed (Fig. 4e,f). These initial movements catalyse a subsequent rearrangement of the actin N-terminal residues that change their interactions with loop2 and helixW, so that the C-terminal end of loop2 is stabilised (Fig. 3i) and actin E4 interacts with the activation loop. The stabilisation of loop2 at its C-terminal end, means that the U50 domain and strut^22^ are pulled towards the actin surface, promoting cleft closure (Fig. 4c). As interactions of the U50 with the actin surface are formed, committing myosin to cleft closure, switch-1 and the P-loop are moved away from switch-2, opening the backdoor, and concomitant reshaping of the nucleotide-binding pocket pushes Pi into the Pi release tunnel (Fig. 4g). Thus, actin catalyses myosin ATPase activity by accelerating cleft closure and Pi dissociation.

Cleft closure is made energetically favourable in the presence of actin, due to the formation of additional interfaces between the myosin U50 and -actin, and the distortions that occur upon binding of primed myosin to actin act to accelerate Pi release. Interactions of Pi with positively-charged residues in the Pi release tunnel^17^ could delay its release into solution and explain why kinetic^7,28^ and single-molecule measurements^14^ suggest that Pi is released after the powerstroke occurs^10^. Cleft closure causes the transducer to twist and the relay helix to straighten, concomitant with lever swing, producing the powerstroke and the postPS structure (Fig. 4h).

In the absence of load, there is tight coupling between cleft closure and lever swing. However, under strain, the lead head of two-headed myosin-5 has been shown to rapidly release Pi^29^ yet adopt a strongly-bound state with a primed lever^30,31^. This is consistent with Pi displacement preceding cleft closure, which commits myosin to lever movement. The activation loop may also have a role here in stabilising the actomyosin interface to decrease detachment under load^32^. The motions of cocking back around the actin axis, and cleft closure are in planes almost orthogonal to that of lever swing, such that neither would be impeded in the presence of load on the lever along the F-actin axis. So rather cleverly, myosin clamps itself onto actin without producing any axial movement. Thus, when the lever tries to swing forward against a restraining force, the axial force doesn’t tend to re-open the cleft. This is akin to how a chameleon climbs up a stick. This feature has important implications for function across all myosin classes.

### Implications for two-headed myosin-5

By overlaying the primed and postPS state actomyosin structures we were able to visualise the lever swing along actin (Fig. 5a-c). If we extend our structures to full lever length (see Methods), the axial working stroke is ∼34 nm, which is consistent with the distance between preferred binding sites on actin^33^ (Fig. 5a). There is a small (4°) right-handed component to the lever swing (Fig. 5b-c) and a small (2.5°) right-handed torsion of the lever around its own axis, such that the lever tips are displaced from one another approximately 7° azimuthally around the actin axis.

**Fig. 5.**
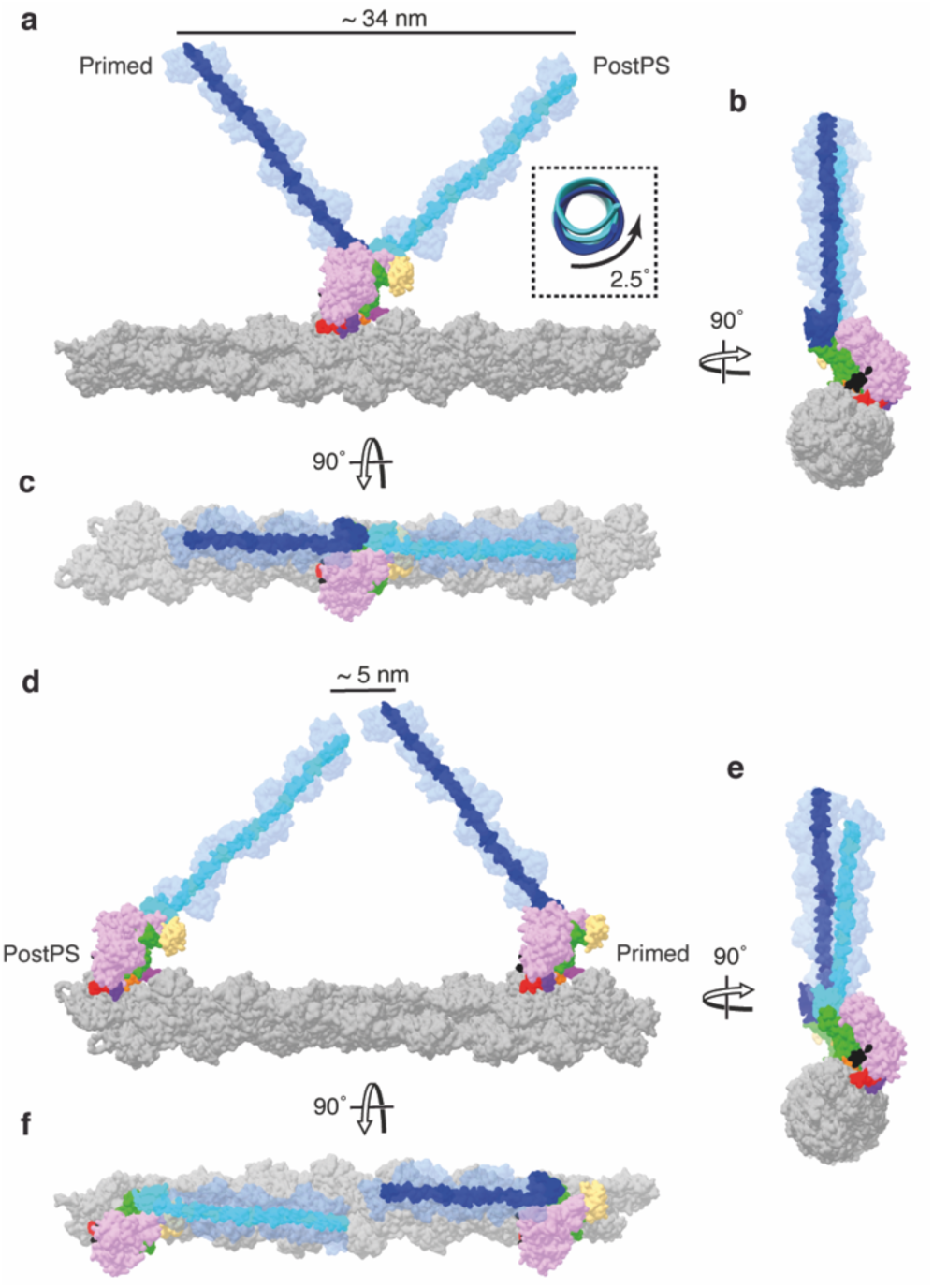
Myosin-5 working stroke and walking on F-actin. (a) Overlay of primed and postPS actomyosin structures with full-length levers, coloured in dark blue and cyan respectively, on actin in side view. A working stroke of approximately 34 nm is seen as well as little rotation of the lever as highlighted. (b) Top and (c) end-on views of the actin filament, show a very small azimuthal displacement of the lever tips (7°). When a postPS and a primed myosin are positioned 13 actin subunits apart, the lever ends meet in a similar position along the actin axis, as observed in (d) side, (e) top and (f) end-on views of the actin filament. Note that this actin filament has a rotation per subunit of -166.6°. Small changes in this value change the relationship of the lever ends in (d-f).

To mimic the walking molecule, we placed a postPS and primed motor, with full-length levers, 13 actin subunits apart along an actin filament, as if they were the leading and trailing head of a myosin-5 double-headed molecule (Fig. 5d-f). The two ends of the levers were slightly displaced from each other azimuthally but met at the same point axially along the filament. This shows that only slight bending of the levers or variation in actin helical symmetry^33^ is needed to unite the heads onto the coiled tail, as is observed by EM^30,34^. During walking, there is thus no need for a forward diffusive search by the detached head.

Together, this means that the myosin-5 motor is able to generate motion very effectively, producing an almost linear motion over a distance that is close to the typical step size along actin.

### Conclusions

By use of time-resolved cryoEM we have captured an actomyosin complex in the primed state and solved its structure to high resolution (4.4 Å). Primed myosin initially binds actin through its lower 50 kDa subdomain. Due to the high conservation in the primed actomyosin interface, the structure of this state is likely conserved across myosin classes and as such, provides a valuable model for understanding the effects of disease-causing myosin mutations. Our time-resolved data show a primed actomyosin structure transitioning to a post-powerstroke structure, directly demonstrating the swinging lever mechanism, and enabling us to propose a mechanism for how actin catalyses it.

## Methods

### Sample preparation

Rabbit skeletal actin in monomeric form (G-Actin) was prepared as previously described ^35^. Polymerisation to F-actin was done by mixing ∼ 300 µM G-Actin with 10 % (v/v) cation exchange buffer (3 mM MgCl_2_, 11 mM EGTA), incubating for 5 min on ice, adding 10 % (v/v) polymerization buffer (120 mM MOPS, 300 mM KCl, 12 mM MgCl_2_, 1 mM EGTA) and incubating the mixture overnight on ice. Mouse myosin-5a head fragment (subfragment 1, S1), coding for amino acids 1– 797 (1 IQ calmodulin-binding motif) and carrying the switch 1 S^217^A mutation, loop 2 DDEK^594-597^ deletion and C-terminal Flag purification tag (Extended Data Fig. 2), was expressed using pVL1392 baculovirus transfer vector and purified as previously described^19^. Disodium ATP was obtained from Roche and ADP was obtained from Sigma Aldrich.

### Kinetic measurements

Transient kinetics of actomyosin ATP hydrolysis were measured by use of an Hitech Scientific stopped-flow with single or double mixing, where appropriate. All stopped-flow experiments were carried out at 20°C with a final buffer concentration of 37.5 mM KAc, 25 mM KCl, 10 mM MOPS (pH 7.0), 2.25 mM MgCl_2_, 0.1 mM EGTA, 0.25 mM DTT in the cell. See Extended Data Fig. 3 for specific method information.

### Time-resolved cryoEM grid preparation

Time-resolved cryoEM experiments were done using a custom-built setup previously described ^21^ with modifications to allow two mixing steps. A photo and schematic of the setup are shown in Extended Data Fig 1. The flow rates for each individual syringe were 2.1 µL/s. In the first mixing step, myosin-5 at 51 µM in 10 mM MOPS, 100 mM KCl, 3 mM MgCl_2_, 0.1 mM EGTA pH 7.0 was mixed 1:1 with 1 mM ATP in reaction buffer (10 mM MOPS, 50 mM KAc, 2 mM MgCl_2_, 0.1 mM EGTA pH 7.0). The myosin-nucleotide mixture at a flowrate of 4.2 µL/s was met by two 2.1 µL/s flows of F-Actin at 25 µM (subunit concentration in reaction buffer) in the flow focussing region of the spray nozzle to create an actin-myosin mixture comprising 13 µM myosin, 13 µM actin, 250 µM ATP, 10 mM MOPS, 38 mM KAc, 25 mM KCl, 2 mM MgCl_2_, 0.1 mM EGTA at pH 7.0, and a total flowrate of 8.4 µL/s. This final mixture was sprayed onto an EM grid.

The average time delay from the first mixing step to the spray nozzle was 2.2 s, given a flowrate of 4.2 µL/s, tube length of 7 cm, inner diameter (I.D.) of 0.38 mm and dead volumes of 1.0 and 0.3 µL for mixer and nozzle, respectively. The spray nozzles used here have been described and characterized previously^20,36^.The nozzle to grid distance at the point of sample application was 1.3 cm and the droplet speed ≥30 m/s, resulting in a time-of-flight for the droplets of less than 1 ms. With a vertical distance of 1.7 cm between spray nozzle and liquid ethane surface and a grid speed of 1.8 m/s, the time-delay was calculated to be 10 ms (10 ± 2 ms). The nozzle was operated in spraying mode with a spray gas pressure of 2 bar.

A longer time-delay of ∼120 ms was obtained by increasing the vertical distance between nozzle and ethane surface to 5.2 cm and pausing the grid after passing the spray. In these experiments, the sample mixture was incubated for an additional ∼100 ms on-grid, before plunging into liquid ethane for vitrification. The total time delay from droplet application to vitrification was 120 ms (122 ± 5 ms), including deceleration, 100 ms pause and acceleration. Otherwise, the conditions for grid preparation were the same as for the 10 ms timepoint.

All grids were prepared at room temperature (∼20 °C) and at >60 % relative humidity in the environmental chamber of the time-resolved EM device. Self-wicking grids were supplied by SPT Labtech and used after glow discharge in a Cressington 208 Carbon coater with glow-discharge unit for 60 s at 0.1 mbar air pressure and 10 mA. Four replicate grids were prepared for each timepoint, 3 of which were taken forward for data collection.

### Data processing and model building

Data were collected on a Titan Krios microscope equipped with a Gatan K2 direct electron detector operated in counting mode. The main data collection and processing parameters are listed in Extended Data Table 1. A schematic overview of the processing pipeline is given in Extended Data Fig. 3. Data from 3 grids were collected for each timepoint. All processing was done using RELION-3.1 ^37^, unless otherwise mentioned. Micrographs were corrected for beam-induced motion using MotionCor2 and CTF estimation was done using GCTF ^38,39^. Actin filaments were manually picked and processed using standard helical processing methods (Extended Data Fig. 3&4) ^40^. After CTF-refinement and Bayesian polishing, all 6 datasets were combined and a helical consensus structure calculated (Extended Data Fig. 4c). Using focussed 3D classification without alignment (non-helical) and a mask that covered the central myosin binding site (Extended Data Fig. 4d), particles were classified into primed actomyosin, postPS actomyosin (Extended Data Fig. 4c, e), free actin and a small fraction of particles left unassigned. The final reconstruction of free actin was obtained by helical refinement. Primed and postPS actomyosin were refined helically and after partial signal subtraction, as single particles (Extended Data Fig. 4f-i). Post-processing was performed in RELION and in DeepEMhancer^41^.

Processing parameters for free myosin-5 are listed in Extended Data 3. Free myosin-5 particles were picked from a subset of micrographs of the 120 ms time-resolved data. Because of thicker ice, free myosin particles were not picked from the 10 ms data. After one round of 2D classification, good particles were used to train a crYOLO model^42^. With the trained model, particles were picked from the entire 120 ms dataset, leading to a final selection of 23930 particles after one round of 2D and one round of 3D classification. The final 3D refinement after Bayesian polishing was done using non-uniform refinement in cryoSPARC^43^.

Homology models were generated using Modeller within Chimera based on the PDB files shown in Extended Data Table 2^44,45^. Model building was done using coot^46^, with subsequent refinement of the nucleotide pocket in ISOLDE, implementing the hydrogen bonding coordination to the phosphates as described in Forgacs *et al.^19^* and TYR119 coordination as described in Pospich *et al.^15^* as harmonic restraints during flexible fitting^25^. Real space refinement was performed using Phenix^47^. To permit elucidation of interactions occurring at the actomyosin interface, we used molecular dynamics simulations. These were performed with the Amber FF14SB forcefield and a GBSA implicit solvent model following the method described in Scarff et al.^48^ Interactions that were observed in at least 50% of the simulations were included in the model. Structures were visualised in Chimera. Videos were generated by use of Chimera, Adobe Aftereffects and Adobe Premiere. For the generation of movie 4, we separated out the motions of cleft closure and powerstroke into a suggested time sequence to produce a model of force generation. To achieve this, a chimeric model of primed and postPS actomyosin was generated (myosin chain numbering: aa1-128 primed, aa129-449 postPS, aa450-507 primed, aa508-632 post, aa633-763 primed).

### Myosin-5 full-length lever model

A 17 actin subunit filament was created by seven superpositions of our three actin subunit model. Full-length levers (to residue 909) were added onto our primed and PostPS actomyosin structures by super-imposing levers from PDB ID 7YV9 chain A, aligned on the converter domain (residues 699-750). Lever swing and azimuthal displacement were measured using the measurement tools in Chimera.

### Data availability

The electron density maps and atomic models for unbound primed myosin-5, primed actomyosin-5 and postPS actomyosin-5 have been deposited into EMDB, with accession codes EMD-19031, EMD-19013 and EMD-19030, and the PDB with accession codes 8RBG, 8R9V and 8RBF, respectively.

The following models were used for comparison purposes in our study, actomyosin-5 rigor structures PDB IDs: 7PLT, 7PLU, 7PLV, 7PLW, 7PLZ and actomyosin-5 strong-ADP structures PDB IDs: 7PM5, 7PM6, 7PM7, 7PM8, 7PM9.

## Acknowledgements

Prof. Howard White, and the late Prof. John Trinick, proposed this experiment some 40 years ago and it has taken until now, with the required improvements in technology, for it to be accomplished. We thank John for his foresight and the mentorship he provided to us. We thank members of the cryoEM community at Leeds for their help and guidance, particularly Glenn Carrington for his advice on movie generation. All EM data were collected at the Astbury Biostructure funded by the University of Leeds and the Wellcome Trust (108466/Z/15/Z; Grant No. 204825/Z/16/Z to C.A.S), and we thank the support scientists for their help in data acquisition. This work was funded by a Biotechnology and Biological Sciences Research Council (BBSRC) grant to S.P.M. (BB/P026397/1), and supported by research grants to H.D.W. from the American Heart Association and to H.D.W. and V.G. from the US National Institutes of Health (NIHR21AR-071675). This work was also funded by the Medical Research Council (Grant No. MR/P018491/1). S.N.M is supported by a School of Molecular and Cellular Biology, University of Leeds, funded PhD studentship. C.A.S is supported by a BHF Jacqueline Murray Coomber Fellowship (FS/20/21/34704).

## Author contributions

H.D.W. and S.P.M. designed the project. B.V. and E.F. produced the myosin mutant constructs. H.D.W., S.P.M., D.P.K. and F.S. aided in design of the time-resolved approach. D.P.K., S.P.M. and H.D.W. performed time-resolved cryoEM grid screening, optimisation and data collection. D.P.K., C.R. and V.G. performed initial data analysis and initial processing of the cryoEM data. E.F., J.A., H.D.W. and D.A.W. performed kinetic experiments and kinetic data analysis. M.S. performed kinetic modelling. S.N.M. and C.A.S. performed cryoEM data refinement and final model building. S.N.M. performed MD simulations. S.N.M. and C.A.S. performed structure validation. S.N.M., C.A.S., S.P.M., P.J.K., D.P.K. and H.D.W. interpreted the data and the model. S.N.M and C.A.S. performed main figure and movie generation. S.N.M., C.A.S., D.P.K., J.A., and S.P.M. produced supplementary figures. C.A.S., S.P.M., P.J.K, D.P.K., S.N.M, and H.D.W. wrote the manuscript. All authors discussed the results and commented on the manuscript.

## Materials & Correspondence

Correspondence and material requests should be addressed to S.P.M, C.A.S, & H.D.W.

---

**Extended Data Table 1:**
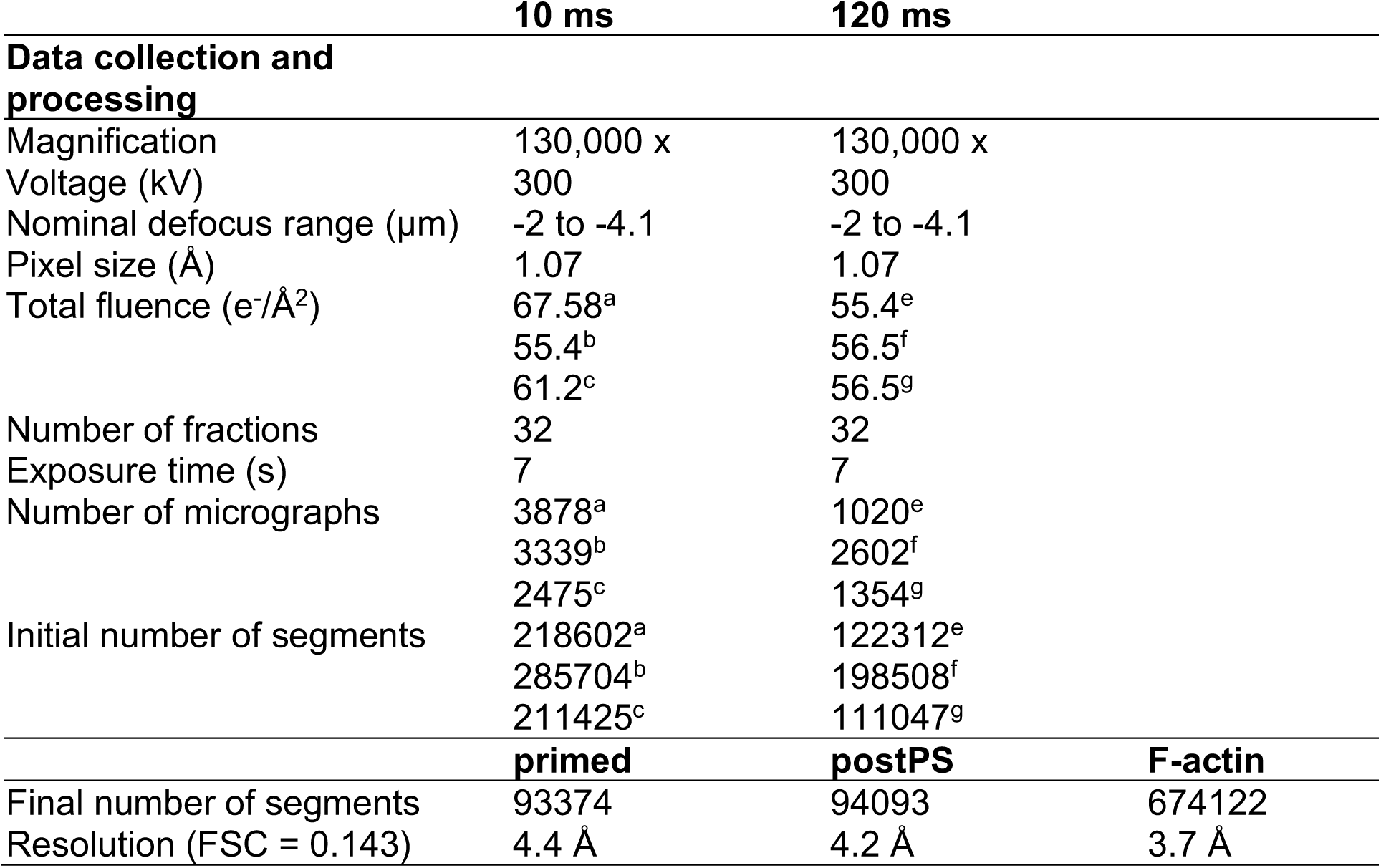
Data collection, processing, model building and refinement statistics for time-resolved EM data, for three collections at 10 ms (^a,b,c^) and 120 ms (^e,f,g^) respectively.

**Extended Data Table 2:** Data collection, processing, model building and refinement statistics for time-resolved EM data.

|  | **Primed**<br>**actomyosin** | **PostPS**<br>**actomyosin** |
| --- | --- | --- |
| **Model Refinement** |  |  |
| Initial Model used | PDB 4ZG4<br>(myosin)<br>PDB 5ONV (actin) | PDB 1W7I<br>(myosin)<br>PDB 5ONV (actin) |
| Map-model correlation (FSC = 0.143) | 4.4 Å | 4.2 Å |
| Map-sharpening B-factor (Å<sup>2</sup>) |  |  |
| RELION post-processing | -119 | -84 |
| **Model composition** |  |  |
| Non-hydrogen atoms | 14852 | 14790 |
| Protein residues | 1857 | 1848 |
| Ligands | 4 | 4 |
| **R.M.S.Z deviations** |  |  |
| Bond lengths (Å) | 0.61 | 0.6 |
| Bond angles (°) | 0.75 | 0.77 |
| **Validation** |  |  |
| MolProbity score | 2.06 | 1.75 |
| Clashscore | 17.12 | 9.73 |
| Poor rotamers (%) | 0.25 | 0.19 |
| **Ramachandran plot** |  |  |
| Favoured (%) | 95.34 | 96.41 |
| Allowed (%) | 4.34 | 3.54 |
| Disallowed (%) | 0.32 | 0.05 |

**Extended Data Table 3:**
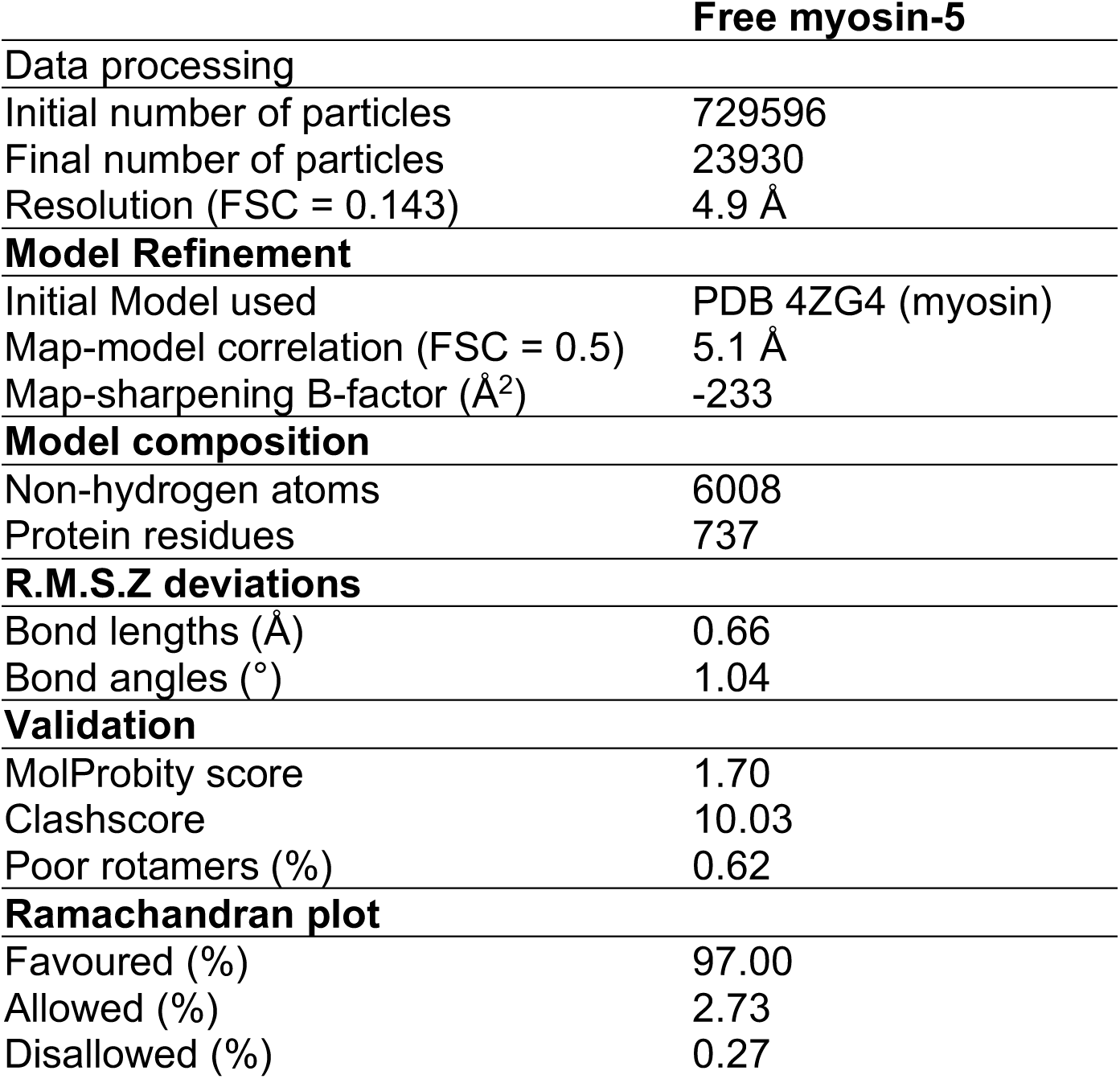
Processing, model building and refinement statistics for unbound primed myosin-5 from 120 ms time-resolved EM data.

## Supporting Information - Swinging lever mechanism of myosin directly demonstrated by time-resolved cryoEM

**Extended Data Fig. 1.**
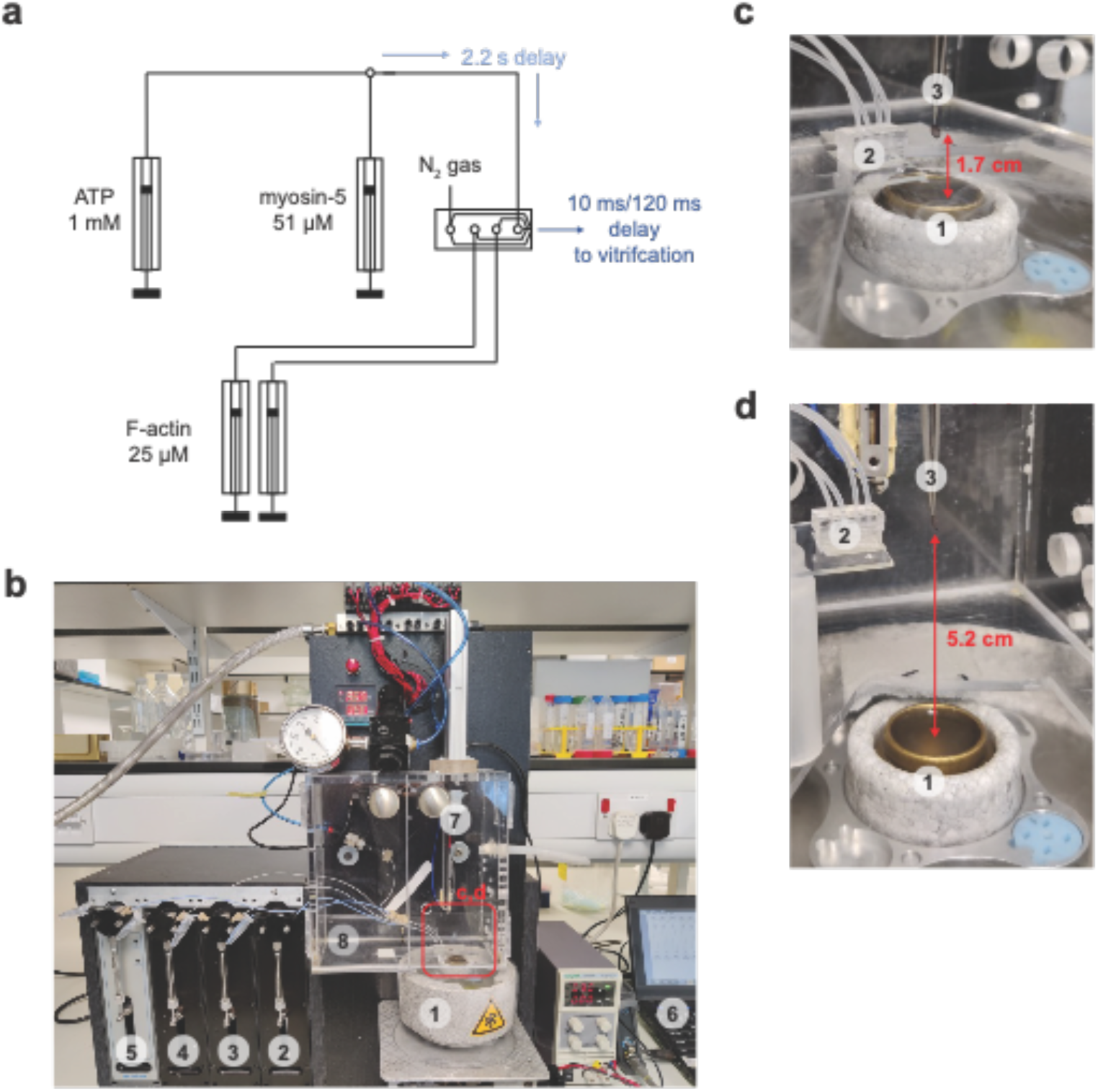
Experimental setup for time-resolved cryoEM. (a) Schematic of experimental setup showing the concentrations of reagents used. (b) Photo of the experimental setup with liquid nitrogen/ethane container (1), syringe pumps (2-5), control PC (6), forceps on plunger (7) and humidity controlled chamber (8). The red box highlights the region around the spray nozzle, magnified views of this region are shown in c-d. (c) Magnified view of ethane cup (1), spray nozzle (2) and grid in sample application position (3) with a short distance for the 10 ms timepoint. (d) Similar to c, except for the larger distance between nozzle and ethane which was used for the 120 ms timepoint.

**Extended Data Fig. 2.**
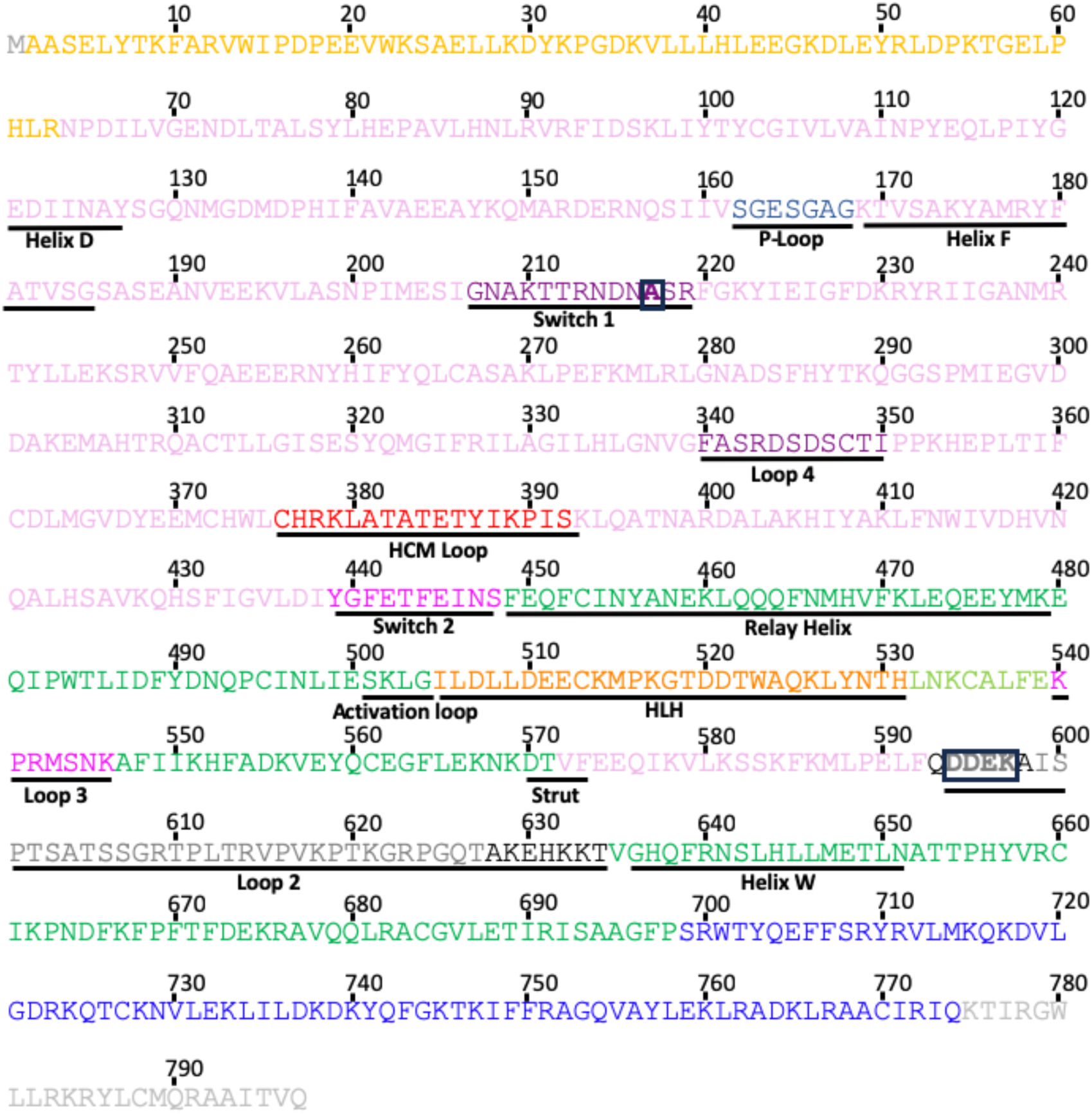
Myosin S1 sequence and domain architecture. Myosin-5 S1 amino acid sequence (myosin heavy chain residues 1-797). Subdomains and regions of interest are colored as in Fig.1-4 and underlined. Gold, N-terminal domain; pink, U50; navy blue, P-loop; purple, switch-1 and loop4; red, HCM loop; magenta, switch-2 and loop3; green, L50; orange, HLH; black/grey, loop2, where residues in black are modelled in our structure and those in grey are not; royal blue, converter (residues 699-750) and modelled region of light chain binding domain; light grey residues 775-797 are the unmodeled region of the light chain binding domain of the construct. The switch 1 S^217^A mutation and loop 2 DDEK^594-597^ deletion are boxed. The construct studied has a FLAG-TAG sequence, DYKDDDDK, C-terminal to the myosin heavy chain sequence stated.

**Extended Data Fig. 3.**
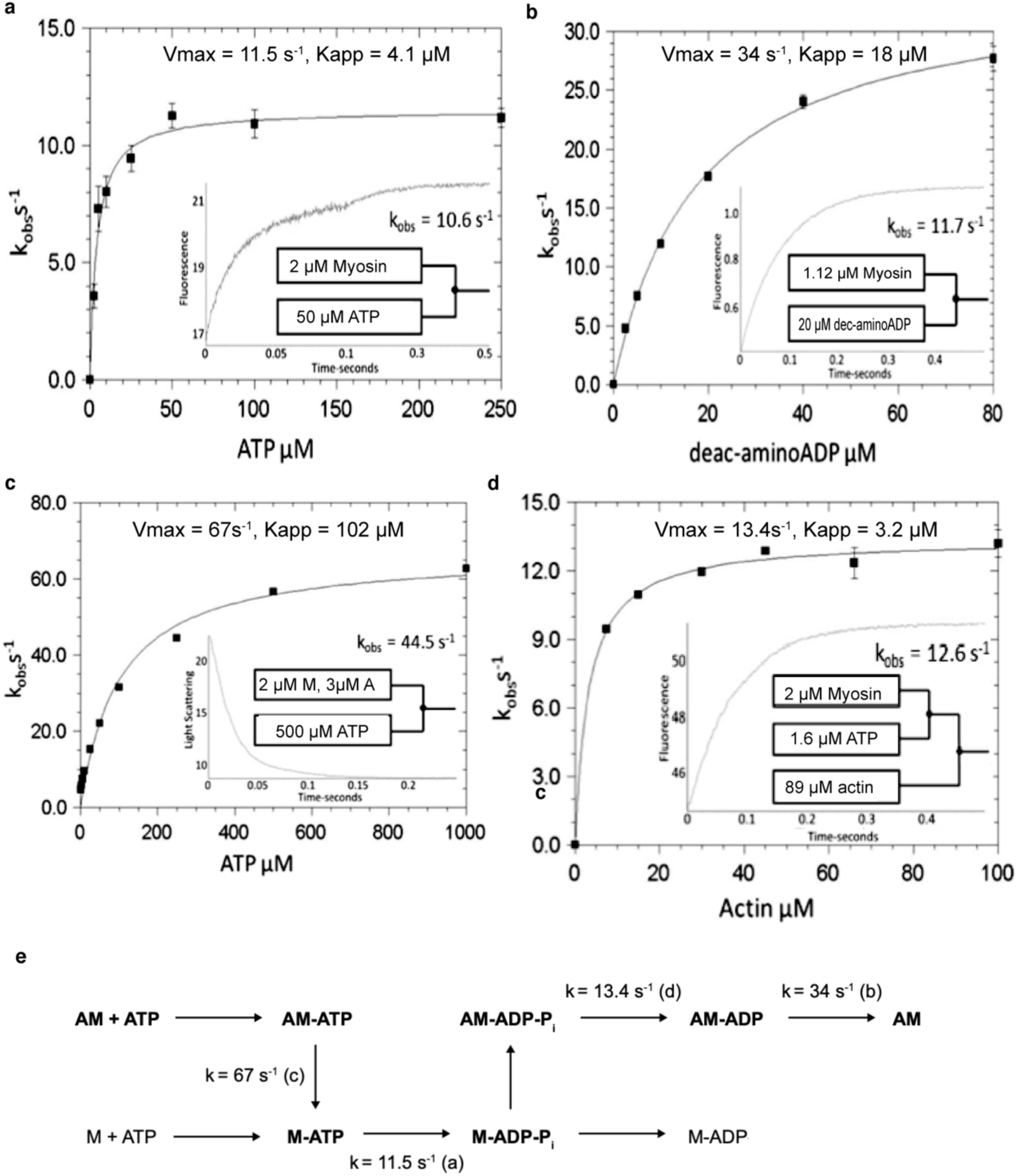
Transient kinetics of mutant actomyosin-5 ATP hydrolysis. ATP hydrolysis measured at 20 °C by single mixing (a-c) or double mixing (d) stopped-flow. Final concentrations in the cell: 37.5 mM KAc, 25 mM KCl, 10 mM MOPS (pH 7.0), 2.25 mM MgCl_2_, 0.1 mM EGTA, 0.25 mM DTT. Representative traces shown in insets with a stopped-flow mixing schematic. **(a)** Myosin ATP hydrolysis measured by intrinsic tryptophan fluorescence using a 320-380 nm bandpass filter with excitation at 295 nm. Final concentrations: 1.0 µM myosin, 1.5 µM calmodulin, and 2.5 - 250 µM ATP. The hyperbolic fit yields V_max_ = 11.5 s^-1^, K_app_ = 4.1 µM. **(b)** ADP dissociation from actomyosin-ADP was measured using a deac-aminoADP chase with a 455 nm long-pass filter and excitation at 430 nm. Final concentrations: 0.14 µM or 0.56 µM myosin and calmodulin, 1.42 µM actin, 5.7 µM ADP, and 2.5 - 80 µM deac-aminoADP. The hyperbolic fit yields V_max_ = 34 s^-1^, K_app_ = 18 µM. **(c)** ATP-induced dissociation of myosin from actin measured by light scattering with a 400 nm long-pass filter and illumination at 432 nm. Final concentrations: 1 µM myosin, 1 µM calmodulin, 1.5 µM actin, and 1 - 1000 µM ATP. The hyperbolic fit yields V_max_ = 67 s^-1^, K_app_ = 102 µM. **(d)** Phosphate dissociation from the actomyosin-ADP-Pi complex, measured by MDCC-PBP with a 455 nm long-pass filter and excitation at 434 nm. 2 µM myosin mixed with 1.6 µM ATP, held in a delay line for 2 s, and then mixed with actin to accelerate P_i_ release. Final concentrations: 0.5 µM myosin, 0.5 µM calmodulin, 0.4 µM ATP, 0 - 100 µM actin, 5 µM MDCC-PBP, 0.1 mM 7- methylguanosine, and 0.01 unit/mL purine nucleoside phosphorylase. The hyperbolic fit yields V_max_ = 13.4 s^-1^, K_app_ = 3.2 µM. (**e**) Kinetic mechanism of mutant actomyosin-5 S1 ATP hydrolysis. Abbreviation: A, actin, M, Myosin-5 S1(1IQ, S217A, ΔDDEK^594-597^), Pi, phosphate. The main actomyosin ATPase pathway is in bold. Parentheses indicate from which experiment the rate and equilibrium constants were obtained.

**Extended Data Fig. 4.**
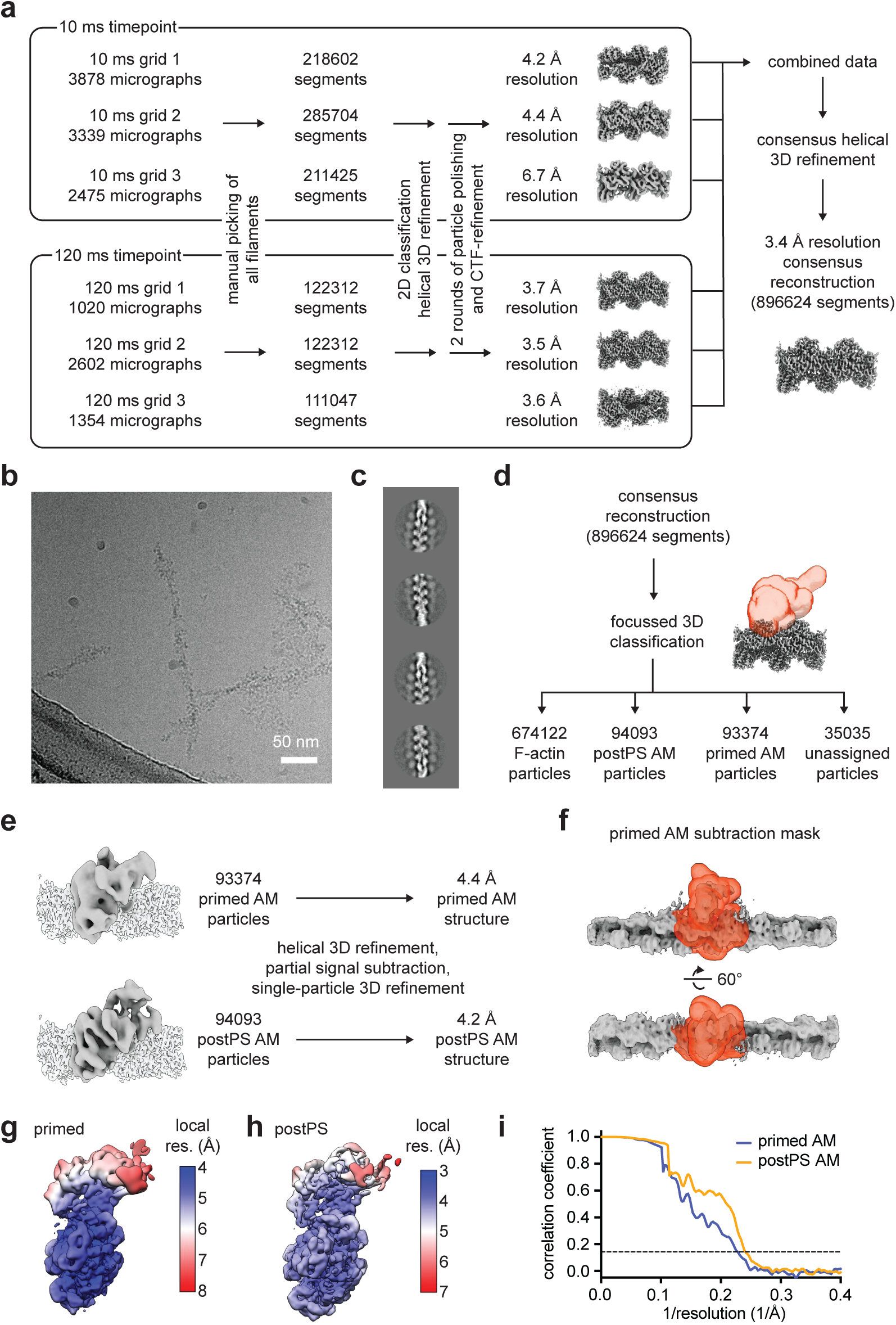
Processing of time-resolved cryoEM data. (a) Flow diagram of the initial image processing of the 10 ms and 120 ms data sets. (b) Micrograph from the 10 ms dataset. (c) 2D classes from the 10 ms timepoint, bound myosin appears as a diffuse density along the actin filament. (d) Result of the focused 3D classification of the combined dataset with a mask covering the myosin binding site (AM: actomyosin). (e) Processing of primed or postPS actomyosin after focused classification. (f) Subtraction mask used for primed actomyosin processing. (g) Final primed actomyosin reconstruction showing local resolution in Å (h) final post PS actomyosin reconstruction showing local resolution in Å (i) Fourier shell correlation curves for prePS (blue) and postPS (yellow) with the 0.143 threshold indicated by a dotted line.

**Extended Data Fig. 5.**
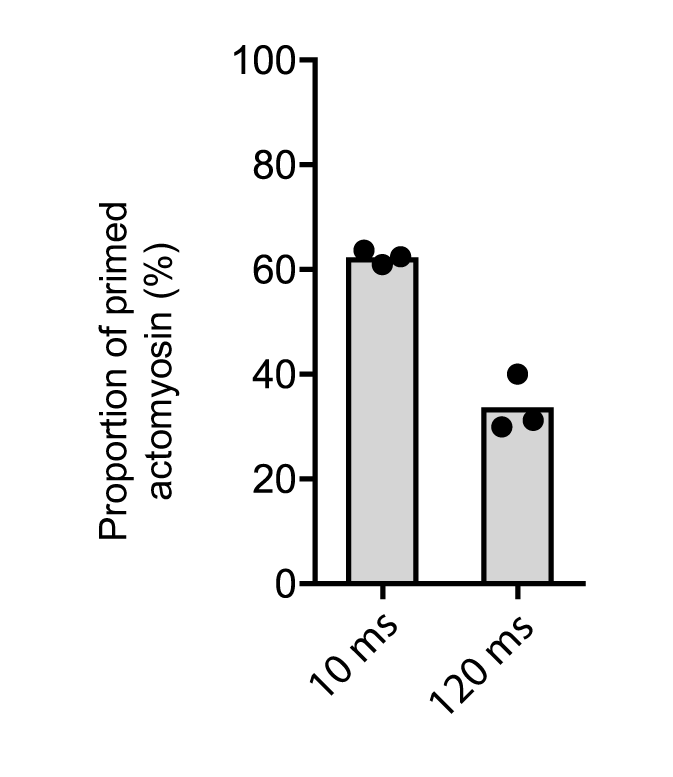
Proportion of primed and postPS states between timepoints and experimental repeats. Proportion of the primed state at 10 ms or 120 ms. Shown is the mean as grey bars and replicates as black points.

**Extended Data Fig. 6.**
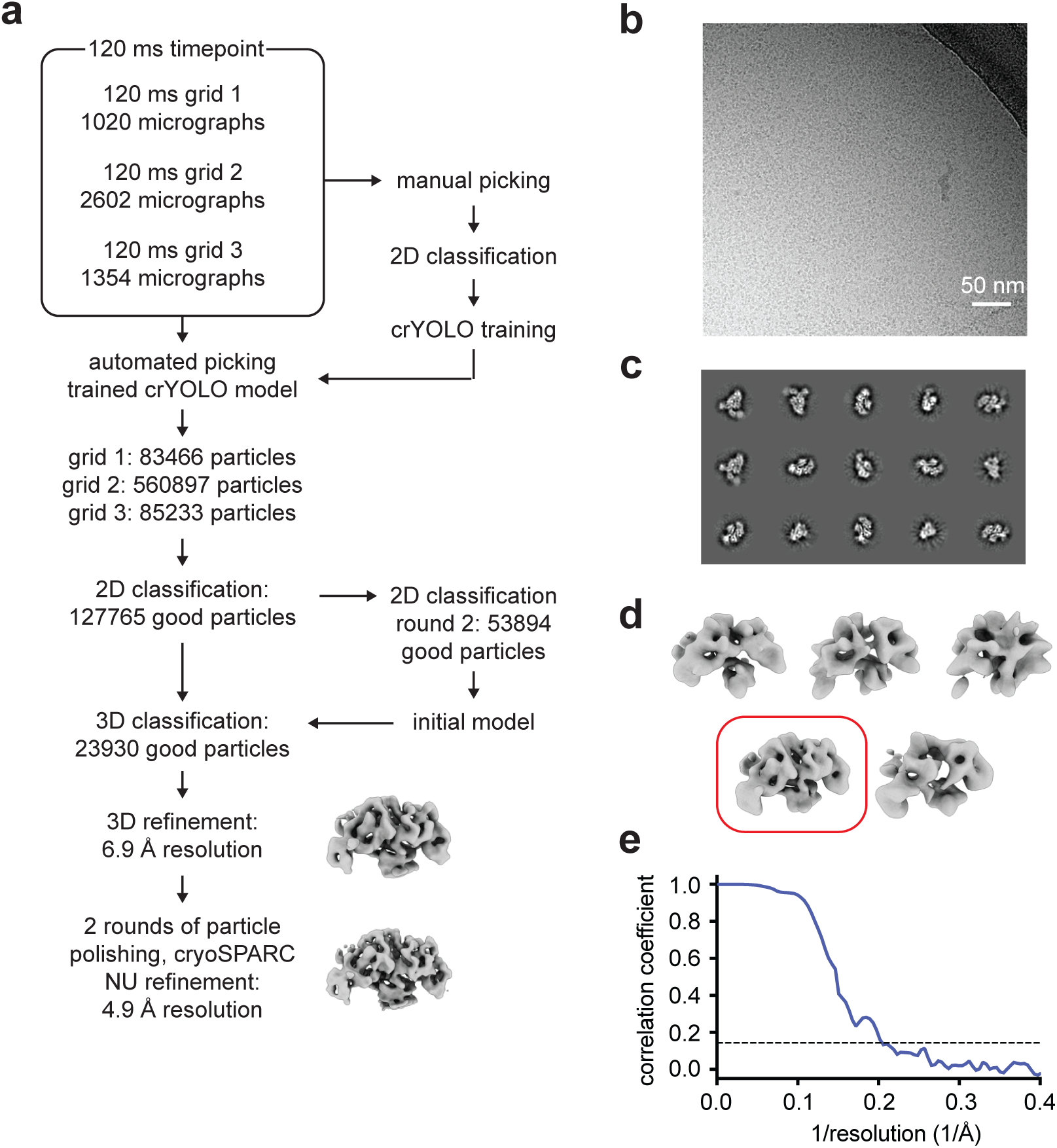
Unbound myosin-5 image processing. (a) Processing pipeline for unbound myosin molecules. (b) Micrograph from the 120 ms time-resolved cryoEM data with a large number of unbound myosin-5 molecules. (c) Representative 2D classes. (d) 3D classification with the selected class highlighted by a red box. (e) Fourier shell correlation curve (blue) with the 0.143 threshold indicated by a dotted line.

**Extended Data Fig. 7.**
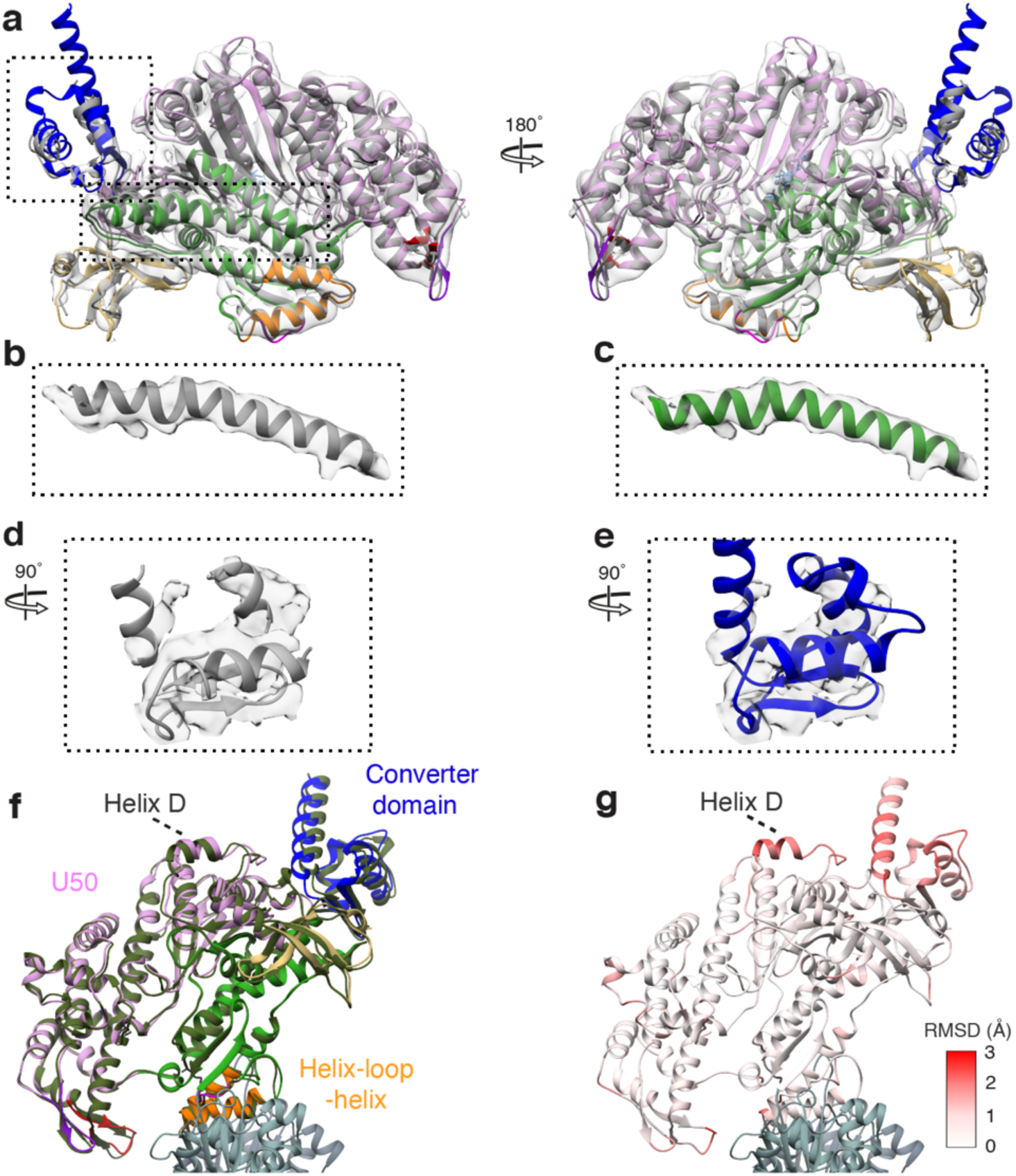
Atomic model of myosin-5a in the primed state. (a) The EM density map of unbound myosin-5 with the crystal structure (PDB ID 4zg4) fitted directly (grey) and after flexible fitting (with subdomains colored, U50 pink, L50 green, N-term gold, converter blue, HLH orange). (b) Relay helix from PDB ID 4zg4 fitted into EM density map for unbound myosin-5 directly and (c) after flexible fitting. (d) Converter domain of PDB ID 4zg4 fitted into EM density map for unbound myosin-5 directly and (e) after flexible fitting. (f) Global superposition of the primed actomyosin-5 (colored as in Fig. 1) and unbound primed myosin-5 (colored olive green) shows a similar structure with no significant changes in domain architecture. (g) RMSD of myosin residues between primed actomyosin and primed myosin.

**Extended Data Fig. 8.**
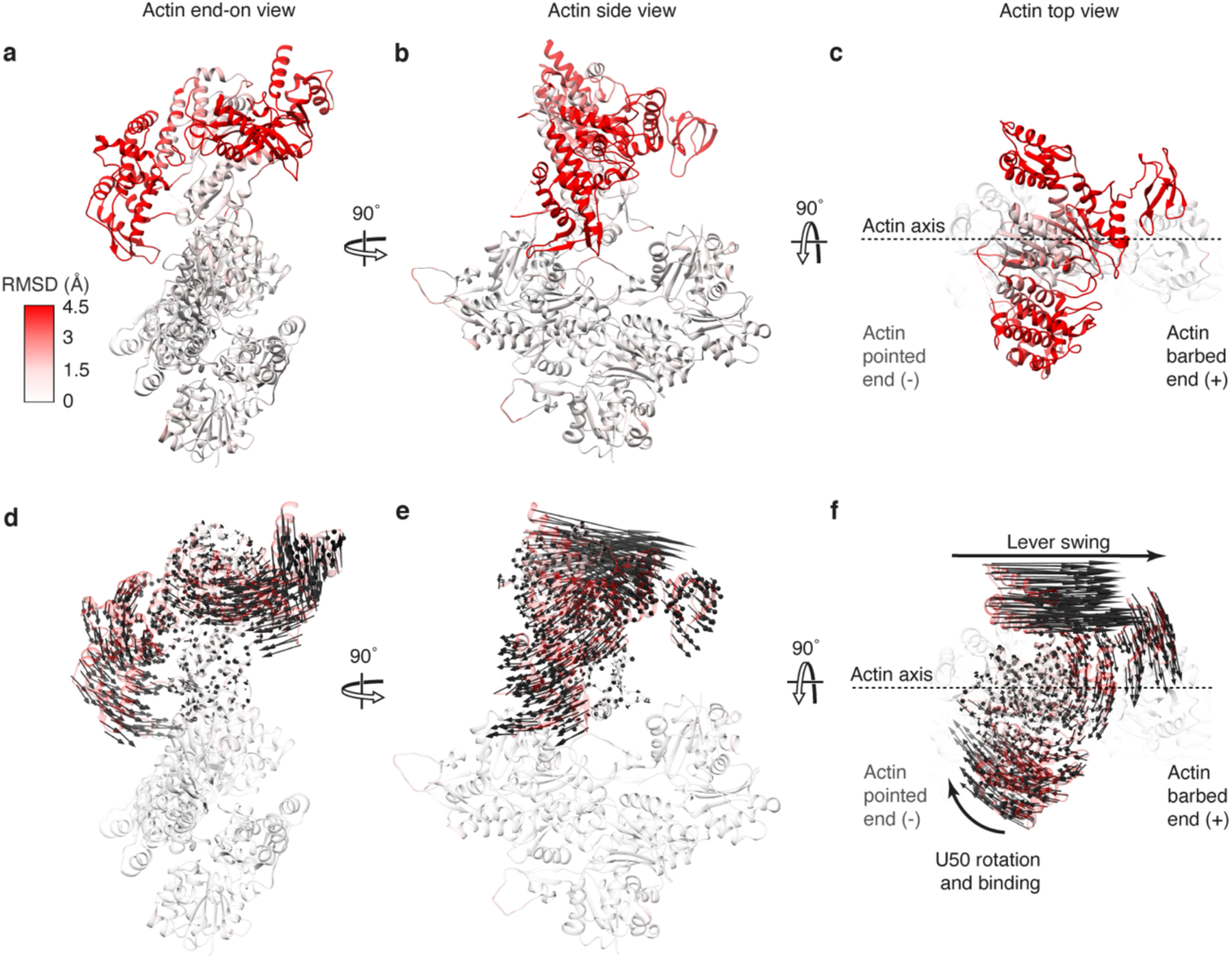
RMSD between primed and PostPS actomyosin structures. Primed actomyosin PDB coloured by RMSD between primed and postPS actomyosin shown in (a) end-on view of F-actin, looking towards the pointed end, (b) view parallel to the actin axis and in (c) top view looking down over the motor domain. (d-f) are the same views as a-c respectively, but with vector arrows (in black) showing displacement in relative Cα positions between primed and postPS actomyosin motor domains.

**Extended Data Fig. 9.**
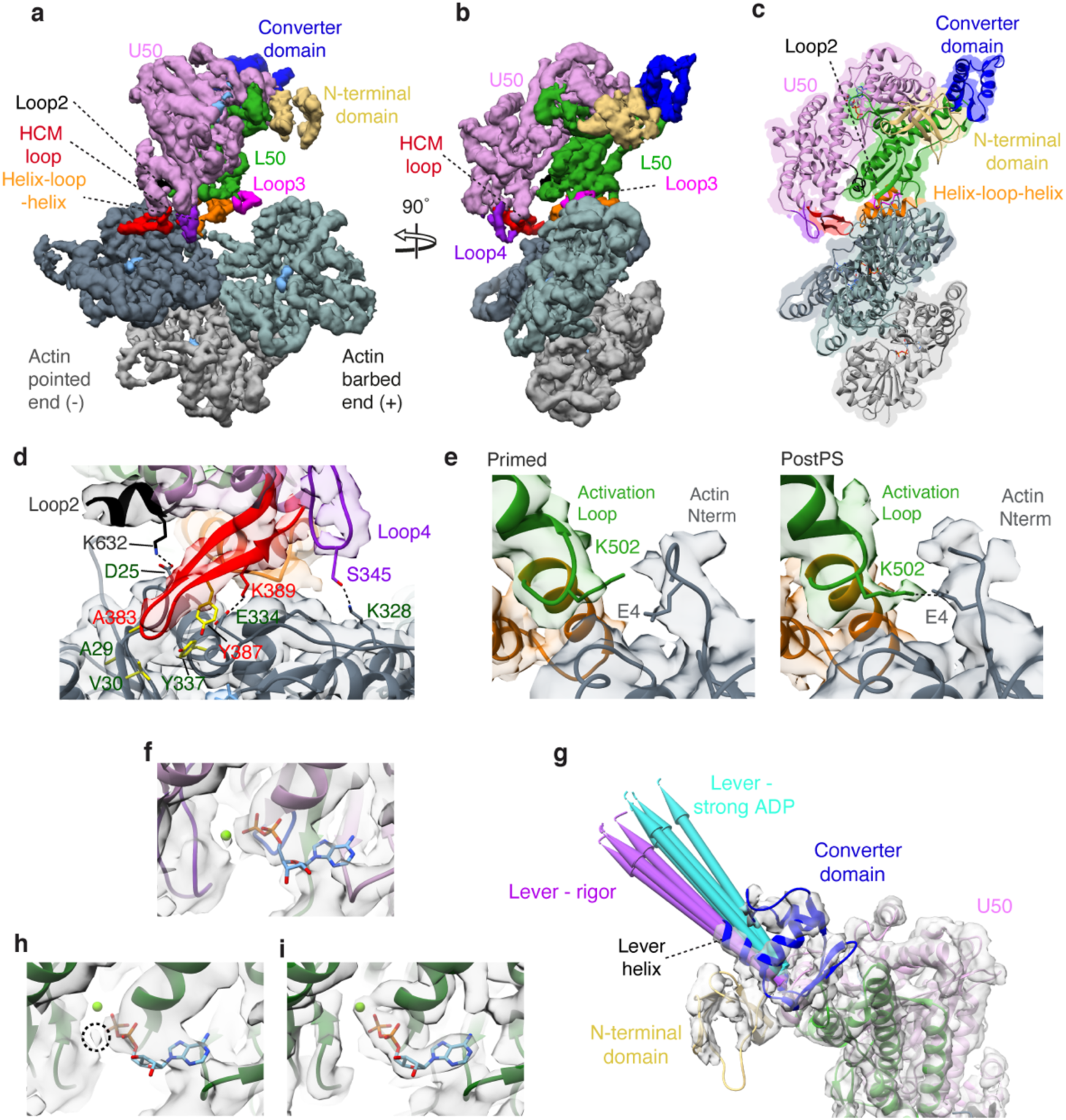
PostPS Actomyosin Structure. CryoEM density map of the postPS actomyosin-5 complex, segmented and colored by myosin subdomains and actin chains as indicated (with central three actin subunits displayed). Map thresholded to show secondary structure (threshold 0.15). Shown in (a) parallel to the actin axis and in (b) in end-on view of F-actin, looking toward the pointed end. (c) Backbone depiction of atomic model of postPS actomyosin-5, fitted into the EM density map, with view as in (b). Actin subunits are shown in slate grey (-end), blue-grey (+end), and light grey. (d) Magnified view of the U50, Loop2, HCM loop and loop4 contacts to actin. Relevant interacting residues are labelled and shown with hydrophobic residues in yellow. (e) Magnified view of the primed actomyosin state and post actomyosin state activation loop-actin N-terminal interaction interface showing formation of hydrogen bond between K502 and E4 in the postPS structure. (f) PostPS nucleotide pocket fit to EM density (map threshold 0.0096), (g) PostPS structure, focused on converter domain. The lever position is more consistent with that observed in previous actomyosin-5 rigor structures (purple pipes, PDB IDs: 7PLT, 7PLU, 7PLV, 7PLW, 7PLZ) than actomyosin-5 strong-ADP structures (turquoise pipes, PDB IDs: 7PM5, 7PM6, 7PM7, 7PM8, 7PM9) (h) Nucleotide pocket of actomyosin structure 7MP5 fitted to our PostPS EM density highlighting unfilled magnesium density with a dashed circle. (i) Nucleotide pocket of actomyosin structure 7MP5 fitted to corresponding density EMDB ID: 13521 (map threshold 0.0197). *DeepEMhancer post-processed map depicted in (a-d, g), and *RELION post-processed map in (e,f,h,i)*.

**Extended Data Fig. 10.**
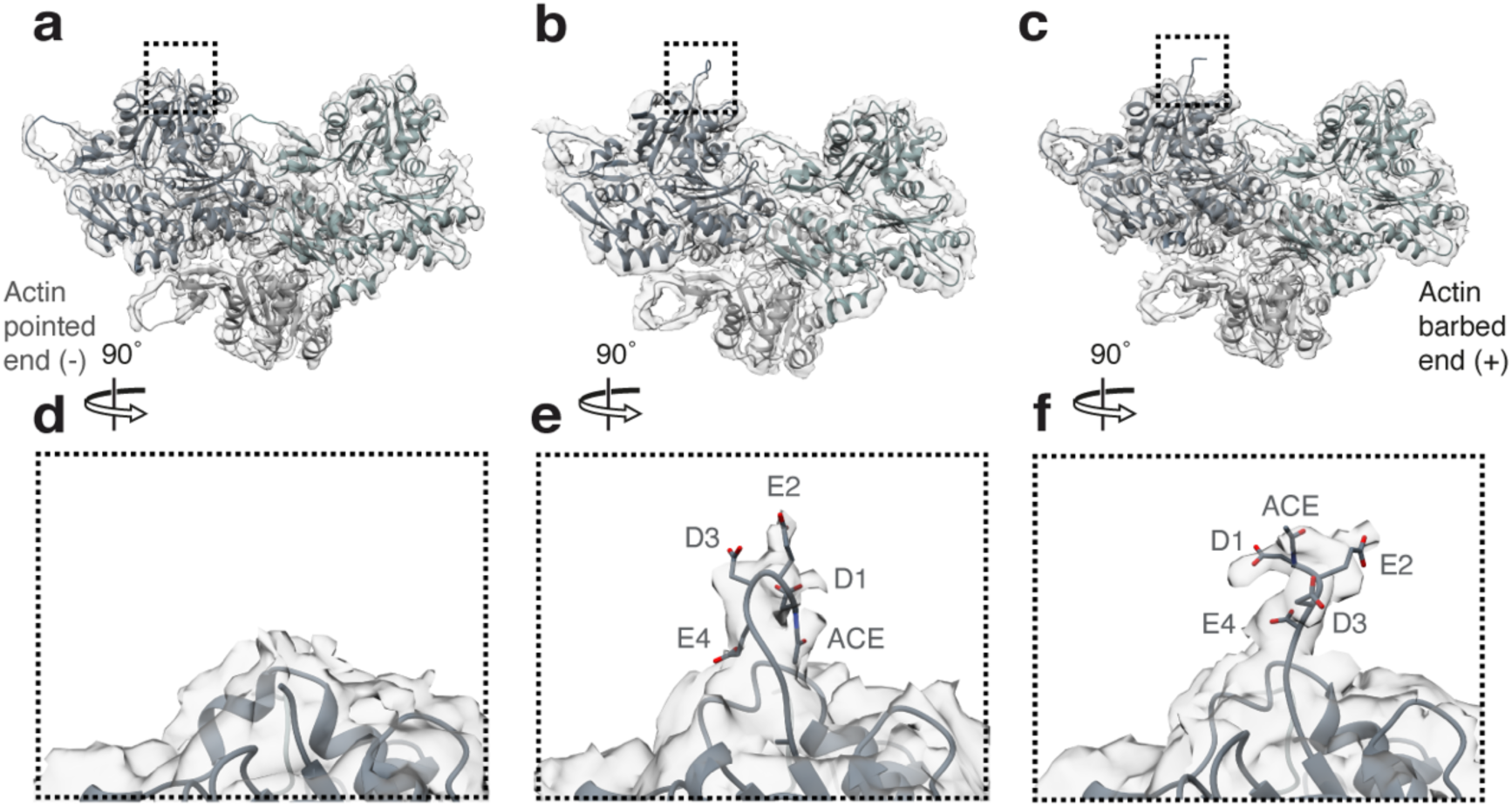
Actin Structure. Actin structure is preserved between (a) actin alone, (b) primed actomyosin, and (c) postPS actomyosin, except at the N-terminus where it becomes ordered when myosin binds. D-loop density also becomes more ordered when associated with myosin. The density observed for the N-terminal four residues of actin is absent in (d) vacant actin and different between (e) primed actomyosin, and (f) postPS actomyosin. ACE = acetyl group of D1.

**Supplementary Video 1. Structure of the primed actomyosin-5 complex.** (Time: 0:00) Transparent cryoEM split map density of the primed actomyosin-5 complex (threshold myosin 0.085, actin 0.2) with backbone depiction of atomic model fitted, rotated 360°. Magnified view of the actomyosin interface fitted to cryoEM map, contacts are made by (0:37) the myosin HLH motif (threshold 0.008), (1:00) loop2 (threshold 0.008), (1:22) actin N-terminus (threshold 0.0065) and (1:52) Loop3 (threshold 0.008).

**Supplementary Video 2. Comparison of myosin structure in the primed actomyosin complex with unbound primed myosin.** (Time: 0:00) CryoEM density map of unbound primed myosin-5 with backbone depiction of atomic model fitted, rotated 360°. (0:32) Depiction of unbound myosin-5 binding F-actin shown looking down the actin axis towards the pointed end. (0:3 s9) Morph from unbound primed myosin-5 to primed actomyosin-5 (aligned on the HLH motif, residues 505-530), highlighting U50 movement and super kinking of the relay helix. (0:50) Reversal of morph. (1:00) Magnified view of the U50 actin binding interface morphing from primed unbound myosin-5 to primed actomyosin-5 highlighting the HCM loop and Loop4’s movement away from the F-actin surface. (1:14) Reversal of morph. (1:33) Magnified view of the nucleotide pocket again morphing from primed unbound myosin-5 to primed actomyosin-5, showing movement of the nucleotide. (1:43) Reversal of morph.

**Supplementary Video 3. Structural changes during the power stroke.** (Time: 0:00) Primed actomyosin cryoEM split map (threshold 0.085) with backbone depiction of atomic model fitted, shown parallel to the actin axis. Morph from Primed actomyosin-5 to postPS actomyosin-5 fitted to relevant cryoEM split maps (threshold primed 0.085 and postPS 0.08) shown (0:13) parallel to the actin axis, (0:42) looking down the actin axis towards the barbed end, (1:09) parallel to the actin axis proximal to the converter, (1:37) looking down the actin axis towards the pointed end and finally (2:13) a top view looking down over the motor domain. (2:46) Magnified view of the actin N-term morphing from primed actomyosin-5 to postPS actomyosin-5 fitted to relevant cryoEM maps (threshold primed 0.0065 postPS 0.0065). (3:34) Magnified view of the U50 actin binding interface morphing from primed actomyosin-5 to postPS actomyosin-5 fitted to relevant cryoEM maps (threshold postPS 0.01). (4:12) Magnified view of the nucleotide pocket morphing from primed actomyosin-5 to postPS actomyosin-5 highlighting the state of the back door.

**Supplementary Video 4. Model of myosin force generation and ATPase activation on F-actin.** Note, here we separated out the motions of cleft closure and powerstroke into a suggested time sequence to produce a model of force generation. To achieve this, a chimeric model of primed and postPS actomyosin was generated (myosin chain numbering: aa1-128 primed, aa129-449 postPS, aa450-507 primed, aa508-632 post, aa633-763 primed). (Time: 0:00) Parallel view of free actin, followed by (0:09) primed myosin initially binding weakly through electrostatic interactions of loop2 with actin subdomain-1. (0:13) This brings the L50 of myosin in close proximity to the actin surface, enabling formation of the stereospecific primed actomyosin state. (0:19) HLH binding enables the actin N-terminal residues 1-4 to interact with HelixW and loop2, resulting in the U50 being cocked back towards the converter domain, which is a rotation around the F-actin axis. (0:25) Rearrangement of N-terminal actin interactions with HelixW and loop2 result in loop2 stabilisation at its C-terminal end, which promotes cleft closure. (0:30) Cleft closure results in the strong binding interface needed for a productive powerstroke. (0:36) Reversal of actin induced motor conformational changes to unbound primed myosin and rotation of view looking down the pointed end of the actin axis. (0:45) Repetition of U50 cocking back, (0:50) actin N-terminal stabilisation resulting in cleft closure and (0:55) productive powerstroke. (1:02) Once again reversal of actin induced motor conformational changes to unbound primed myosin and rotation of view looking down the barbed end of the actin axis. (1:14) Repetition of U50 cocking back highlighting the closed state of the back door and ADP movement. (1:20) Actin N-terminal stabilisation resulting in cleft closure, (1:24) opening the back door allowing Pi release and (1:29) a productive powerstroke.

